# Nuclear NPR1 Modulates Salicylic Acid/Jasmonic Acid Crosstalk via Transcriptional Control of WRKY Transcription Factors

**DOI:** 10.64898/2026.10.20.760389

**Authors:** Niels Aerts, Lotte Caarls, Tessa W. Visscher, Berend Groutars, Renée Kapteijn, Marcel C. Van Verk, Corné M.J. Pieterse, Saskia C.M. Van Wees

**Affiliations:** Plant-Microbe Interactions, Department of Biology, Science4Life, Utrecht University, P.O. Box 800.56, 3408 TB Utrecht, The Netherlands

**Keywords:** Crosstalk, jasmonic acid, NPR1, ORA59, salicylic acid, WRKY

## Abstract

The plant hormones salicylic acid (SA) and jasmonic acid (JA) are major regulators of plant defense. They can antagonize each other’s activity, enabling an optimized defense response. The SA master regulator NPR1 is a key player in this SA/JA crosstalk, although the mechanisms of NPR1-mediated SA/JA crosstalk remain only partly understood, especially regarding the ethylene-coregulated ERF branch of the JA pathway. Here, we investigated the function of NPR1 in SA/JA crosstalk using several Arabidopsis *npr1* mutant variants. Assays using the *NPR1-HBD* and *npr1-nls* lines, expressing only cytosolic NPR1, revealed that NPR1 has a nuclear role in SA/JA crosstalk. RNA-seq analysis of lines expressing NPR1 mutated in residues Cys^82^ or Cys^216^, in which repression of the JA pathway was impaired while SA pathway induction remained mostly intact, identified 32 SA-responsive genes with reduced expression compared to Col-0, including four WRKY transcription factor genes. Of these, WRKY53 and WRKY70 could suppress expression of the JA marker gene *PDF1.2* induced by ORA59, the master regulator of the ERF branch of the JA pathway, in a protoplast system. Additional SA-inducible WRKYs (WRKY50, WRKY54, WRKY63, and WRKY67) similarly suppressed ORA59-induced *PDF1.2* expression, implicating WRKY63 and WRKY67 for the first time in SA/JA crosstalk and confirming earlier reports on the role of the other WRKY TFs. WRKY53 and WRKY67 were further shown to reduce accumulation of the ORA59 protein in protoplasts. Thus, our study indicated that NPR1 can regulate SA/JA crosstalk in the nucleus via transcriptional activation of genes encoding specific JA-repressing WRKY transcription factors.

**MAIN CONCLUSION:** NPR1 has a nuclear function in regulating salicylic acid/jasmonic acid crosstalk by activating specific WRKY transcription factors that suppress the jasmonic acid pathway, likely in part by reducing ORA59 accumulation.

## INTRODUCTION

The plant immune system is activated upon recognition of specific molecules derived from pathogens, insects or damaged host cells (Dodds and Rathjen 2010). A successful defense response stops infection and signals systemic tissues to become primed for enhanced defense against future attacks (Gao et al. 2015). Within this immune signaling network, plant hormones serve as key regulators of specific immune network sectors (Pieterse et al. 2012; Tsuda et al. 2009). Salicylic acid (SA) and jasmonic acid (JA) are major defense hormones that regulate the localized response to attacks and the establishment of systemic resistance in distant, still healthy tissues. Generally, SA is essential in defenses against biotrophic pathogens, which feed on living host cells, while the JA pathway is more effective against herbivorous insects and necrotrophic pathogens, which kill host tissue and feed on the contents (Glazebrook 2005).

Activation of the SA or JA pathway initiates extensive transcriptional reprogramming, leading to the activation of a distinct set of pathogenesis-related (*PR*) genes by both hormones (Hickman et al. 2019; Hickman et al. 2017; Van Loon et al. 2006). The SA pathway is primarily mediated by the transcriptional regulator NPR1, which is required for the activation of numerous SA-responsive genes, including the SA marker gene *PR1*, and for conferring SA-dependent disease resistance (Cao et al. 1994; Delaney et al. 1995; Yun et al. 2024; Wang et al. 2006; Powers et al. 2024; Zavaliev and Dong 2024). NPR1 interacts with TGA transcription factors (TFs) to drive downstream gene expression, including that of *PR1* and various WRKY TF genes that fine-tune and enhance downstream immune responses (Wang et al. 2006; Blanco et al. 2009; Pandey and Somssich 2009; Powers et al. 2024; Yun et al. 2024). Activation of JA-responsive genes relies on the JA-induced degradation of JAZ repressor proteins, which, under basal conditions, suppress the activity of JA-responsive TFs (Thines et al. 2007; Chini et al. 2007). Degradation of JAZs is triggered by JA-Ile, the bioactive form of JA, which facilitates the interaction of JAZs with the SCF^COI1^ ubiquitin ligase complex, freeing the previously repressed TFs (Pauwels and Goossens 2011). Two branches are distinguished in JA-dependent defense signaling. The first is coregulated by ethylene and controlled by ERF TFs such as ORA59 (Pré et al. 2008), while the second is coregulated by ABA and controlled by MYC TFs such as MYC2 (Fernández-Calvo et al. 2011). These branches are referred to as the ERF branch and the MYC branch, respectively. The ERF branch is generally effective against necrotrophic pathogens, while the MYC branch is mostly directed against herbivorous insects and is also activated upon wounding (Pieterse et al. 2012). On the transcriptional level, ERF branch activity is usually assessed by the expression of the marker gene *PDF1.2*, a target gene of ORA59, whereas MYC branch activaty is usually determined by the expression of the marker gene *VSP2*, a target of MYC2 (Zarei et al. 2011; Pieterse et al. 2012; Dombrecht et al. 2007).

Hormonal signaling pathways involved in plant defense form a complex network that operates at multiple levels of regulation (Aerts et al. 2021). This network includes numerous antagonistic and synergistic interactions between pathways, collectively known as hormonal crosstalk (Pieterse et al. 2012; Robert-Seilaniantz et al. 2011). Hormonal crosstalk in the context of plant defense ensures an optimal response given the nature of the attacker, abiotic conditions, developmental stage and time of day (Atkinson and Urwin 2012; Lu et al. 2017; Nobori and Tsuda 2019). Well-known examples of defense-related hormonal crosstalk include the antagonism between the SA and JA pathways and between the MYC and ERF branches of the JA pathway (Pieterse et al. 2012; Aerts et al. 2021). Several studies have helped to clarify the molecular mechanisms underlying SA-mediated suppression of the JA response (from hereon: SA/JA crosstalk). In the model plant *Arabidopsis thaliana* (from hereon: Arabidopsis), SA has been shown to significantly affect the expression pattern of 69% of the more than 3500 MeJA-responsive genes (Hickman et al. 2019), highlighting the extensive impact of SA on JA signaling. SA further targets the JA pathway downstream of JA biosynthesis and of SCF^COI1^ through multiple mechanisms, including direct repression of JA-induced transcription and degradation of the positive ERF branch regulator ORA59 (Leon-Reyes et al. 2010b; Van der Does et al. 2013; He et al. 2017; Nomoto et al. 2021). In addition, several SA-controlled transcriptional (co-)regulators that suppress JA-dependent gene expression have been identified (Caarls et al. 2015). These include the TGA TFs TGA2, TGA5, and TGA6 (Leon-Reyes et al. 2010a; Zander et al. 2014) as well as the WRKY TFs WRKY41, WRKY46, WRKY50, WRKY51, WRKY53, WRKY62, and WRKY70 (Li et al. 2004; Hu et al. 2012; Mao et al. 2007; Higashi et al. 2008; Yan et al. 2018; Gao et al. 2011). By contrast, SA-induced EAR-motif-containing transcriptional repressors from the ERF family do not appear to play a role in SA/JA crosstalk (Caarls et al. 2017).

NPR1 is recognized as a key transcriptional coregulator in SA/JA crosstalk. It is required for SA-mediated suppression of the JA marker genes *VSP2*, *PDF1.2*, and *LOX2* (Spoel et al. 2003; Nomoto et al. 2021), and plays a crucial role in controlling SA-mediated suppression of JA-induced resistance against herbivorous insects and necrotrophic pathogens in Arabidopsis (Leon-Reyes et al. 2009; Spoel et al. 2007). The importance of NPR1 in SA/JA crosstalk extends to tomato, rice, and *Nicotiana attenuata* (Yuan et al. 2007; El Oirdi et al. 2011; Rayapuram and Baldwin 2007), indicating a conserved role for NPR1 as an SA-activated modulator of the JA pathway. However, the mechanisms by which NPR1 suppresses the JA pathway are not fully elucidated yet.

Studies on how SA modulates NPR1 may shed light on possible mechanisms of NPR1-mediated SA/JA crosstalk. SA-mediated regulation of NPR1 and subsequent activation of SA-responsive genes have been studied extensively (Chen et al. 2021; Zavaliev and Dong 2024), and the crystal structure of NPR1 is solved (Kumar et al. 2022). The NPR1 protein contains several conserved domains essential for its function as a transcriptional coactivator. The ankyrin repeat domain mediates interaction with TGA TFs, which is required for *PR* gene activation (Zhang et al. 1999; Després et al. 2003; Zhou et al. 2000; Kumar et al. 2022), while the BTB domain enables NPR1 oligomerization (Kumar et al. 2022) and interacts with TGA2 to counteract the effects of TGA2’s N-terminal repression domain, thereby promoting downstream transcription (Boyle et al. 2009). Under non-inducing conditions, the BTB domain acts as an auto-inhibitory domain that masks the NPR1 C-terminal transactivation domain (Wu et al. 2012). This C-terminal transactivation domain is also the site where SA binds to NPR1. Binding of SA disrupts the interaction between the C-terminal domain and the BTB domain, and induces the docking of the C-terminal domain onto the ankyrin repeats 3 and 4, thereby converting NPR1 into an activated transcriptional coactivator (Wu et al. 2012; Manohar et al. 2015; Rochon et al. 2006; Kumar et al. 2022). The transcriptional activity mediated by NPR1 is further boosted by the formation of SA-induced nuclear NPR1 condensates (Powers et al. 2024).

The localization of NPR1 also affects its function. Under non-inducing conditions, intramolecular disulfide bonds between cysteine residues promote the formation of oligomers, thereby sequestering NPR1 in the cytosol. Meanwhile, any of the few monomers that still enter the nucleus are subjected to proteasomal degradation (Spoel et al. 2009). Changing either Cys^82^ or Cys^216^ to alanine reduces NPR1 oligomerization while increasing nuclear localization, showing the importance of the cysteine residues in regulating the oligomer/monomer status of NPR1 (Mou et al. 2003; Kumar et al. 2022). In addition, S-nitrosylation of Cys^156^ of NPR1 increases oligomerization (Tada et al. 2008). Accumulation of SA causes a redox change in the cell, after which thioredoxins TRX-h3 and TRX-h5 reduce the cysteine residues, breaking the intramolecular disulfide bonds and releasing NPR1 monomers that can move to the nucleus via nuclear pore proteins (Mou et al. 2003; Tada et al. 2008; Cheng et al. 2009). In the nucleus, dimerization of NPR1 is important for activating SA-related transcription, for example, by bridging two DNA-bound TGA3 dimers (Kumar et al. 2022).

While nuclear localization is essential for NPR1’s function as a transcriptional coactivator, its necessity in the regulation of SA/JA crosstalk remains unclear. For example, while the interaction with TGA TFs is important for activation of SA-responsive genes, one study showed that the TGA binding site in the *PDF1.2* promoter is not required for SA/JA crosstalk acting on this gene (Spoel et al. 2003). In the same study, Spoel et al. (2003), suggested that cytosolic localization of NPR1 is sufficient to suppress the ERF branch marker gene *PDF1.2*, based on a DEX-inducible *NPR1-HBD* line where NPR1 is located in the cytosol. Further supporting the cytosolic role, overexpression of OsNPR1 in rice suppressed JA-responsive genes and reduced defense against an insect herbivore, while Cys^76^ and Cys^216^-mutated NPR1, which was nuclear localized, was unable to suppress JA-responsive genes under basal conditions, lowering the resistance level against insect herbivores (Yuan et al. 2007). Another possible mechanism for cytosolic regulation of crosstalk by NPR1 comes from a study where it was shown that NPR1 can form cytosolic SA-induced condensates containing stress- and immunity-related proteins, which undergo ubiquitination and degradation (Zavaliev et al. 2020). However, these bodies were not enriched for components related to JA signaling (Zavaliev et al. 2020), suggesting that these cytosolic NPR1 condensates are not involved in NPR1-mediated SA/JA crosstalk. In support of a nuclear role, one study found that NPR1 can suppress the MYC branch of JA signaling by binding MYC2 at the same position as the mediator subunit MED25 (Nomoto et al. 2021). This binding blocks the MYC2-MED25 interaction, which is an essential interaction for full MYC branch activation (Çevik et al. 2012; Chen et al. 2012). While this discovery largely clarified the role of NPR1 in suppressing the MYC branch, the mechanisms by which NPR1 suppresses the ERF branch remain to be fully elucidated. Altogether, this illustrates that the role of NPR1 localization in SA/JA crosstalk is not clear-cut and suggests that NPR1 can regulate SA/JA crosstalk via multiple mechanisms, possibly depending on the context.

In this study, we used several transgenic lines with altered NPR1 to investigate whether nuclear localization of NPR1 is required for SA/JA crosstalk, focusing mainly on the lesser-studied ERF branch of the JA pathway. We found that SA/JA crosstalk is impaired in the *NPR1-HBD* and *npr1-nls* mutants, which carry variants of NPR1 that are retained in the cytosol, suggesting that nuclear localization is required for NPR1’s role as a transcriptional repressor of the JA pathway. Follow-up experiments revealed that two lines overexpressing NPR1 with mutations at either Cys^82^ or Cys^216^ were impaired in SA-mediated suppression of JA marker gene expression, but unaffected in SA-induced *PR1* expression. This distinction separates NPR1’s role as a transcriptional activator in the SA response from its transcriptional repressor role in SA/JA crosstalk. Further analysis of affected genes in these lines uncovered several WRKY TFs as novel players in NPR1-regulated SA/JA crosstalk. Experiments in protoplasts suggested that they act on the ERF branch downstream of the ORA59 TF protein, partly via reducing ORA59 protein levels, resulting in reduced *PDF1.2* expression.

## MATERIALS AND METHODS

### Plant material and growth conditions

In all experiments, Arabidopsis Col-0 was used as wild type and it was the background accession for all mutant and transgenic lines studied. The mutants and overexpression lines used were generated by previous studies and were: *35S_pro_*:*NPR1-HBD npr1-1* (from hereon: *NPR1-HBD*), *35S_pro_*:*npr1-nls-GFP npr1-1* (from hereon: *npr1-nls*) (Kinkema et al. 2000), *npr1-1* (Cao et al. 1994), *35S_pro_:NPR1^C82A^-GFP npr1-1* (from hereon: *C82A*), and *35S_pro_:NPR1^C216A^-GFP npr1-1* (from hereon: *C216A*) (Mou et al. 2003).

For experiments with mature plants, sowing and plant growth was performed as follows. Arabidopsis seeds were stratified in the dark for 48 h at 4°C in 0.1% agar and then sown in boxes on river sand that had been autoclaved twice for 45 min with a 24-h interval. The boxes were then put in larger trays with water at the bottom and a closed transparent lid to ensure 100% relative humidity. Two weeks after germination, seedlings were transferred to 60-mL pots containing a sand/potting soil mixture (5:12 v/v) that had been autoclaved twice for 45 min with a 24-h interval. Plants were cultivated in trays on a wetted cloth in a growth chamber with a 10-h day and 14-h night cycle at 70% relative humidity at 21°C. Plants were watered two times a week by soaking the cloth with water, and one additional time each week they were supplied with a modified half-strength Hoagland solution that also contained 10 µM sequestrene (Fe-EDDHA, Royal Brinkman’s-Gravenzande, the Netherlands).

For experiments with seedlings, seeds were first surface sterilized by mixing 100 mL of commercial bleach (Manutan B.V., Den Dolder, the Netherlands) with 3.2 mL 37% HCl in a beaker in a desiccator, after which the desiccator was closed immediately and the seeds were incubated in the desiccator for 4 h. For protoplast experiments, 50 µL of sterilized seeds were sown directly onto 120×120-mm square Petri dishes that contained 70 mL of ½ Murashige and Skoog (MS) medium with 2% (w/v) sucrose, buffered with MES (pH=5.7) and solidified with 1% plant agar. For hormone treatment experiments, sterilized seeds were sown in 120×120-mm square Petri dishes on an autoclaved, fluid-permeable nylon membrane (Sefar Nitex 03-100/44) that lay on top of 50 mL of solidified ½ MS medium (same as for protoplasts except 0.8% agar). Seeds were sown in two rows on the plates, making sure that all plates had five seeds of each line. For both types of experiments, the seeds were stratified on the plates in the dark at 4°C for 3 d and then incubated lying down (protoplast experiments) or upright (hormone experiments) in a growth chamber at 21°C with a 10-h light/14-h dark cycle at a light intensity of 75 μmol m^-2^ s^-1^ (protoplast experiments) or 100 μmol m^-2^ s^-1^ (hormone experiments).

### Chemical treatments

For hormone treatment of mature plants, 5-week-old plants were treated with SA and/or MeJA by dipping the leaves into a solution containing 0.015% (v/v) Silwet L77 (Van Meeuwen Chemicals BV, Weesp, the Netherlands; nowadays this compound is known as CoatOSil 77 (Momentive, New York, NY, USA)) and either 0.1 mM MeJA (Serva, Brunschwig Chemie, Amsterdam, the Netherlands), or 1 mM SA (Mallinckrodt Baker, Deventer, the Netherlands) and 0.1 mM MeJA. For mock treatments, plants were dipped into a solution containing 0.015% (v/v) Silwet L77 and 0.1% ethanol, as ethanol was the solvent for the 1000x MeJA stock. Leaf number 8 (counting from the oldest) was harvested for RNA isolation 5 h and 24 h after treatment.

For hormone treatments of seedlings, nylon membranes with 12-day-old seedlings previously grown on ½ MS agar plates were transferred to new plates containing the same ½ MS agar medium, but containing either 0.02 mM MeJA (diluted from a 5000x stock in ethanol), 0.5 mM SA + 0.02 mM MeJA, or a mock treatment (½ MS agar medium with 0.02% ethanol). Plants were incubated upright in a growth chamber for 48 h under the conditions described above. Harvesting was done by putting one seedling of a genotype from each of five plates in a 2-mL tube (Greiner Bio-One, Kremsmünster, Austria) containing two glass beads, and immediately snap-freezing it in liquid nitrogen. A total of five replicates were harvested, where one biological replicate consisted of five plants from five different plates.

### RNA extraction, qRT-PCR and RNA-seq

For qRT-PCR analysis, RNA was extracted as described for vegetative tissues by Oñate-Sánchez and Vicente-Carbajosa (2008). RNA was pretreated with DNAse I (Thermo Fischer Scientific, Waltham, MA, USA) to remove genomic DNA. RevertAid H minus Reverse Transcriptase (Thermo Fisher Scientific) was used to convert DNA-free total RNA into cDNA. PCR reactions were performed in optical 384-well plates with a ViiA 7 realtime PCR system (Applied Biosystems, Waltham, MA, USA), using SYBR^®^ Green to monitor the synthesis of double-stranded DNA. The primers used to analyze expression by qRT-PCR are described in Table S1 (File S1). A standard thermal profile was used: 50°C for 2 min, 95°C for 10 min, 40 cycles of 95°C for 15 s and 60°C for 1 min. Amplicon dissociation curves were recorded after cycle 40 by heating from 60°C to 95°C with a ramp speed of 1.0°C min^-1^. Expression was calculated as -ΔCt relative to the reference gene *AT1G13320* (Czechowski et al. 2005) and normalized to the mean - ΔCt value of mock-treated Col-0.

For RNA-seq, developmental leaf 8 was harvested from three individual mock-treated wild-type Col-0 plants or SA + MeJA-treated Col-0, *npr1-1, C82A*, and *C216A* plants, 5 h after treatment. RNA was extracted using the RNeasy Plant Mini Kit (Qiagen, Hilden, Germany) according to the manufacturer’s instructions, including the optional on-column DNAse treatment step. RNA quality was assessed on the 2100 Bioanalyzer (Agilent, Santa Clara, CA, USA) using the RNA 6000 Nano Labchip Kit (Agilent). Library preparation was done using the TruSeq mRNA Stranded Sample Prep kit (Illumina, San Diego, CA, USA) and samples were sequenced on the NextSeq 500 platform (Illumina) with read lengths of 75 bases.

### Analysis of RNA-seq results

Reads were aligned to the Arabidopsis genome (TAIR v10) using TOPHAT v2.0.4 (Trapnell et al. 2009) using the following settings: ‘transcriptome-mismatches 3’, ‘N 3’, ‘bowtie1’, ‘no-novel-juncs’, ‘genome-read-mismatches 3’, ‘p 6’, ‘read-mismatches 3’, ‘G’, ‘min-intron-length 40’, ‘max-intron-length 2000’. Aligned reads were summarized over annotated gene models (TAIR v10) using HTseq-count version 0.5.3p9 with default settings. Sample counts were depth-adjusted and differential expression was determined using the DESeq package with default settings (Anders and Huber 2010). Genes with a corrected *P* value (*P* adjusted) of ≤ 0.05 were called as differentially expressed. Expression of the *npr1-1* allele was estimated by multiplying the total number of *NPR1* counts with the number of counts containing the C1000T point mutation of the *npr1-1* allele divided by the number of counts containing either the C1000T point mutation or the wild-type C at position 1000. The rest of the counts were assigned to the wild-type *NPR1* allele (for Col-0) or the C>A *NPR1* allele (for *C82A* and *C216A*). Principal component analysis was performed on log_2_-transformed depth-adjusted reads using the stats package (v4.5.0) in R (v4.1.2).

### Construction of plasmids for protoplast expression assays

Plasmids used for transfection into protoplasts for transient expression assays were assembled with Golden Gate cloning using the Moclo kit (Weber et al. 2011; Werner et al. 2012). For creation of level 0 plasmids, a 1.2 kb fragment of the *PDF1.2* promoter (directly upstream of the translation start site) was amplified from gDNA of Col-0, the CDSs of *ORA59* and of several WRKY TF genes were amplified from cDNA, and the *LucR* CDS was amplified from the pBS-35S-Rluc plasmid. Primers used are listed in Table S2 (File S1). The primers introduced a restriction site for BbsI and a specific sequence of 4 bp that would become the overhang needed to clone the PCR product into the correct vector. For some constructs an internal restriction site needed to be removed from the CDS. This was done by PCR-amplification of two fragments around this site using primers that introduced a nucleotide change so that the restriction site was removed. Simultaneously, a BbsI restriction site was introduced to be able to re-assemble the fragments in the level 0 reaction (File S1 - Table S2). The PCR product was purified after gel electrophoresis using the Illustra GFX PCR DNA and Gel Band Purification Kit (Merck, Darmstadt, Germany). Several small DNA fragments were generated by annealing two custom-designed single stranded oligonucleotides (Integrated DNA Technologies, Coralville, IA, USA). This was done by mixing 25 mL of each (100 mM) in a 1.5-mL tube, followed by heating at 95°C in tap water and allowing the tube to slowly cool down to room temperature. The resulting annealed fragment contained the required overhang of 4 bp on each side for the ligation in a level-0 reaction (File S1 - Table S2).

The PCR products and annealed fragments were cloned into the appropriate vector (Table S2) using a standard Golden Gate reaction. For this, 2 mL of 10x buffer G, 0.2 mL of 100 mM ATP, 1 mL of 10 U mL^-1^ BbsI (Thermo Fisher Scientific), 0.4 mL of 5 U mL^-1^ T4 DNA ligase (Thermo Fisher Scientific), 100 ng of vector DNA, and PCR/annealing product in an approximate molar ratio of 1:2 were mixed in a total volume of 20 mL. The mix was then incubated for 3 cycles of 10 min at 37°C and 10 min at 22°C, which, after the final cycle, was followed by 10 min at 37°C and 20 min at 65°C. The product was transformed into *Escherichia coli* DH5a, after which the amplified plasmid was isolated using the GenElute^TM^ Plasmid Miniprep Kit (Sigma-Aldrich, St. Louis, MO, USA) and the correct insertion of the PCR fragment(s) was verified using Sanger sequencing.

The level 1 reactions followed a similar procedure as the level 0 reactions, with the exception of using different vectors (File S1 - Table S3) and the restriction enzyme BsaI (Thermo Fisher Scientific). Correct plasmid assembly was confirmed through a restriction reaction. Subsequently, various combinations of level 1 vectors were cloned into distinct level 2 vectors (File S1 - Table S4) using a Golden Gate reaction, employing the restriction enzyme BbsI. In this particular reaction, instead of 3 cycles, 12 cycles were conducted, consisting of 10 minutes at 37°C followed by 10 minutes at 22°C. The final construct was confirmed by a restriction reaction.

### Preparation of protoplasts for transient expression assays

Transient expression assays in protoplasts were conducted using a custom protocol based on Yoo et al. (2007) and Mathur and Koncz (1998). Rosettes of 3-to-4-week-old Arabidopsis plants grown on plate (see ‘Plant material and growth conditions’) were cut off from their roots using a razor blade and immediately submerged in 12.5 mL of protoplasting enzyme solution consisting of Cellulase R10 (1.5% (w/v); Duchefa Biochemie, Haarlem, the Netherlands), 0.4% (w/v) Macerozyme R10 (Duchefa Biochemie), 0.4 M Mannitol, 20 mM KCl, 20 mM MES (pH=5.7), 10 mM CaCl_2_ and 0.1% BSA in a 60-mm Petri dish. The enzyme solution was prepared by heating everything except CaCl_2_ and BSA to 55°C for 10 min and adding the last two components after cooling the solution to room temperature. After the rosettes were submerged, they were cut into fine strips (±1 mm) using a razor blade. Next, they were put under vacuum for 2x 5 min using the Savant DNA 110 SpeedVac Concentrator with drying rate set to ‘low’. The suspension was briefly swirled in between. From this point onward the protoplast suspension was handled with extreme care, avoiding excessive movement. After the vacuum infiltration, the plate was wrapped in aluminum foil and incubated at 28°C for 3.5 h, with gentle swirling every 30 min. Subsequently, the protoplast suspension was gently pipetted through a 70-mm filter into a 50-mL tube (Greiner Bio-One). The tube and filter were previously prepared by pipetting 5 mL of W5 solution (0.2 mM MES (pH=5.7), 154 mM NaCl, 125 mM CaCl_2_, and 5 mM KCl) through the filter into the tube. After filtering the protoplast suspension, another 5 mL of W5 was added to the plate, the plate was gently swirled, and the suspension was pipetted through the filter into the tube. Next, the tube was centrifuged at 200 *g* for 2 min in the swing out Multifuge X3R Refrigerated Centrifuge (Thermo Fisher Scientific) at room temperature. Acceleration was set at 5 (out of 9) and brake at 3 to minimize mechanical stress. The supernatant was removed, and the protoplasts were resuspended in 10 mL of W5 and incubated on ice for 30 min. The suspension was centrifuged at the same settings and in case the supernatant was not clear, the addition of W5, incubation on ice and centrifugation was repeated. After removal of clear supernatant the protoplasts were resuspended in 5 mL of MMG solution (4 mM MES (pH=5.7), 0.4 M Mannitol, 15 mM MgCl_2_). The protoplast density was determined in a hemocytometer, and after centrifuging at the same settings and removing the supernatant, the protoplasts were resuspended in MMG to a density of 1×10^6^ cells mL^-1^, after which they were immediately transfected.

### Transfection of protoplasts with a plasmid

Protoplast transfection was done in 12-well cell culture plates with relatively high hydrophobicity (Greiner Bio-One, Item No. 665102). Per transfection a protoplast suspension droplet of 100 mL was pipetted in the middle of each well and 110 mL of PEG solution (40% (w/v) PEG4000 (Fluka; Sigma-Aldrich), 100 mM CaCl_2_, 200 mM mannitol) was added around the edge of the well to avoid mixing with the protoplasts. Approximately 20 μg of plasmid DNA (concentration range 4-12 μg μL^-1^) was added to the protoplast suspension in the well and gently mixed while pipetting. After 5 min of incubation, the plate was swirled to allow mixing between the PEG solution and the protoplast suspension. After 10 min of incubation in a tilted position, 440 μL of W5 was added to stop transfection and the content of each well was transferred to a 2-mL tube (Greiner Bio-One). The tubes were centrifuged for 2 min at 1000 *g* at the ‘soft’ setting in the fixed-rotor 5424 R centrifuge (Eppendorf, Hamburg, Germany). After careful removal of the supernatant, the protoplasts were resuspended in 700 μL of WI solution (4 mM MES (pH=5.7), 20 mM KCl, 0.5 M Mannitol) and immediately transferred to 6-well cell culture plates (Greiner Bio-One) that already contained 500 μL of WI solution. The plates were incubated at 21°C in the dark for approximately 20 h.

Before harvesting, the transfection efficiency was estimated under the ZEISS Axio Zoom.V16 microscope (Carl Zeiss, Oberkochen, Germany) based on EYFP fluorescence in transfected protoplasts. When the efficiency was at least 20%, the protoplasts were harvested. Experiments where multiple samples had less than 20% transfection efficiency were discarded. For harvesting, the protoplast suspension was transferred to 2-mL tubes and centrifuged at maximum speed for approximately 10 s in the 5424 R centrifuge at the ‘soft’ setting. Supernatant was removed and the tubes with the protoplast pellet were snap-frozen in liquid nitrogen and then stored at -80°C until further processing.

### Measuring transient expression in transfected protoplasts

For GUS, LucR and LucF measurements the protoplast pellet was resuspended in 50 mL of lysis buffer (2.5 mM Tris-phosphate (pH=7.8), 1 mM DTT (1,4-dithiotreitol), 2 mM DACTAA (*trans*-1,2-diaminocyclohexane-N,N,N′,N′-tetraacetic acid), 10% (v/v) glycerol, 1% (v/v) Triton X-100). Protoplasts were ruptured by vortexing for 2 s, briefly spinning in a tabletop centrifuge at 2000 *g*, and vortexing again for 1 s, after which the suspension was incubated on ice for 30 min. After warming up to room temperature the suspension was centrifuged at 1000 *g* for 2 min in the 5424 R centrifuge.

For the GUS assay, 10 mL of supernatant of the centrifuged protoplast lysate (from one well of transfected protoplasts per biological replicate) was mixed with 100 mL of freshly prepared MUG substrate (10 mM Tris-HCl (pH=8.0), 2 mM MgCl_2_, 1 mM 4-MUG (4-Methylumbelliferyl-b-D-glucuronide)) and incubated at room temperature for 90 min. Next, 990 µl of 0.2 M Na_2_CO_3_ was added and mixed by pipetting up and down three times to stop the reaction. The solution was then divided into 5x 200 mL in a clear F-bottom 96-well plate (Greiner Bio-One), serving as five technical replicates. Fluorescence of the solution was then determined on the Synergy™ HTX Multi-Mode Microplate Reader (Agilent), with excitation at 360 nm and emission at 460 nm. Measurements were done three times to minimize impact of technical variation. Therefore, the GUS measurement for one sample was the mean of three measurements of five wells.

For the LucF and LucR measurements the remaining lysate (from one well of transfected protoplasts per biological replicate, corresponding to the GUS measurements) was divided into 3x 10 mL in non-transparent, white F-bottom 96-well plates (Greiner Bio-One). Luciferase measurements were done on the Promega^TM^ GloMax® Plate Reader. For experiments where only LucF was measured the Luciferase Assay System kit (Promega, Madison, WI, USA) was used, and for dual luciferase measurements the Dual-Luciferase® Reporter Assay System kit (Promega) was used. First, the luciferin (single luciferase) or the LARII and Stop&Glo solutions (dual luciferase) were prepared from the kits according to the manufacturer’s instructions. Next, one (single luciferase) or two (dual luciferase) injectors of the machine were primed with the appropriate solutions, after which LucF or LucF and LucR measurements were done per well with delay set at 2 s and integration time at 10 s. The measurements consisted of injection of 25 mL of luciferin followed by measurement of LucF (single luciferase), or sequential injection of 50 ml of the LARII solution followed by measurement of LucF, and then injection of 50 mL of the Stop&Glo solution followed by measurement of LucR (dual luciferase). The measurement of one sample for both LucF and LucR was the mean of the measurements of three wells in which the lysate of one sample was divided.

### Data analysis tools and statistical analysis

Data analysis and statistics were done using R version 4.1.2. For RNA-seq data, differential expression was calculated using the DESeq package (Anders and Huber 2010). For all other data, normal distribution and equality of variances were verified using Shapiro–Wilk and Levene tests, respectively. Differences between conditions were then determined using ANOVA followed by Tukey’s HSD test or Dunnett’s test. Visualization was done using the ggplot2 v3.3.5 and gplots v3.3.1 packages. Colors were picked using the RColorBrewer v1.1–2 package and via http://www.ColorBrewer.org (Cynthia A. Brewer, accessed March/April 2026). The statistics used and the number of biological replicates in each analysis is stated in the legend of the relevant figure. The definition of biological replicates and the technical replicates for each experiment are given in the corresponding section of Materials and methods.

## RESULTS

### Nuclear NPR1 regulates SA/JA crosstalk in seedlings and mature plants

To determine whether nuclear or cytosolic NPR1 is required for SA/JA crosstalk, we modified an experiment by Spoel et al. (2003), who discovered that cytosolic NPR1 could still mediate SA/JA crosstalk on the ERF branch marker gene *PDF1.2*. We grew plants for 12 days on ½ MS agar plates, then transferred them to fresh ½ MS agar plates containing MeJA or SA + MeJA, and incubated them for 48 h. To assess the importance of nuclear NPR1 localization, we used two lines impaired in nuclear NPR1 localization. The line *35S_pro_*:*NPR1-HBD npr1-1* (*NPR1-HBD*) has NPR1 fused to the glucocorticoid receptor hormone binding domain, ensuring cytosolic localization in the absence of dexamethasone (Kinkema et al. 2000). This line was previously also used by Spoel et al. (2003). The other line, *35S_pro_*:*npr1-nls-GFP npr1-1* (*npr1-nls*), has a mutated nuclear localization signal, causing exclusive cytosolic NPR1 localization (Kinkema et al. 2000). As a positive control, we used *npr1-1*, which has a non-functional version of NPR1, and is also the background of the *NPR1-HBD* and *npr1-nls* lines. As expected, SA + MeJA treatment induced the SA marker gene *PR1* in wild-type Col-0, but not in *npr1-1* or the lines with cytosolic NPR1 (File S1 - Fig. S1a). Importantly, while MeJA-induced *PDF1.2* expression was similar between Col-0 and the two lines with cytosolic NPR1, expression after combined SA + MeJA treatment led to significantly lower expression in Col-0 than in the lines with cytosolic NPR1 (Fig. 1a). In this combined treatment, *PDF1.2* expression was reduced in Col-0 compared to the single MeJA treatment, whereas expression levels in *npr1-1*, *NPR1-HBD* and *npr1-nls* were comparable between the SA + MeJA and MeJA treatments. This suggests that nuclear localization of NPR1 is required for SA/JA crosstalk under the experimental conditions used. We repeated the experiment with soil-grown 5-week-old plants, using a rosette hormone dip method of rosettes that is common in our lab to investigate hormone responses in mature plants (Hickman et al. 2017; Aerts et al. 2024; Hickman et al. 2019). The results mirrored those in seedlings (Fig. 1b; File S1 - Fig. S1b-d), suggesting that nuclear NPR1 is required for SA/JA crosstalk in both life stages. Notably, crosstalk regulation of the MYC branch marker gene *VSP2* showed a less pronounced dependence on NPR1 nuclear localization than that of *PDF1.2*, although there was still a significant difference in crosstalk in most NPR1 mutants compared to Col-0 (File S1 - Fig. S1d, e).

**Fig. 1.**
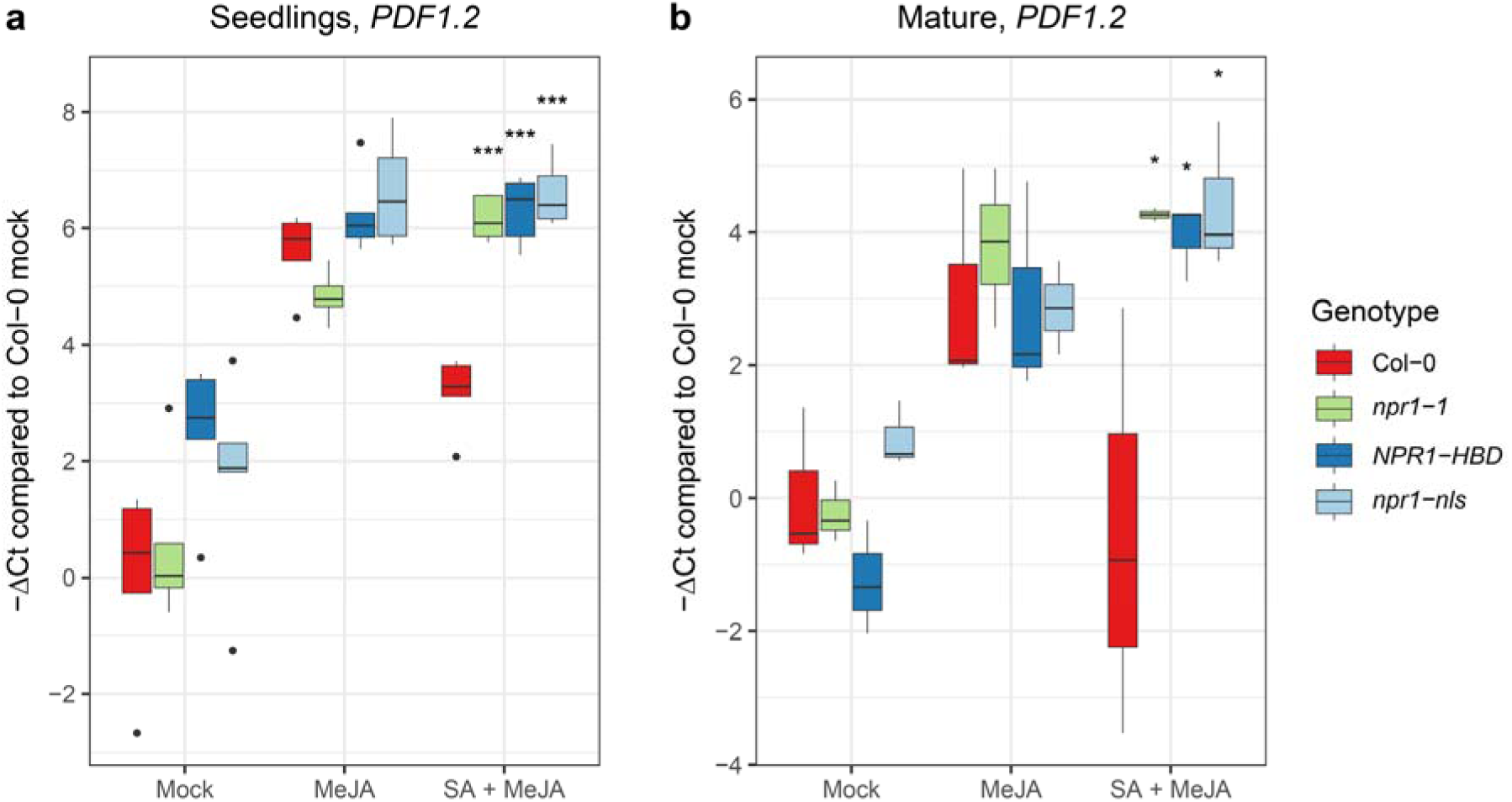
Effects of mutations affecting NPR1 localization on SA/JA crosstalk. qRT-PCR expression analysis of JA marker gene *PDF1.2* after hormone treatments in various localization-related NPR1 mutants. (**a**) Gene expression measured in 12-day-old seedlings on plates. Plants were treated for 48 h with MeJA (0.02 mM) and/or SA (0.5 mM) or mock. (**b**) Gene expression measured in 5-week-old plants in pots. Rosettes were dipped in hormone solution (0.1 mM MeJA and/or 1 mM SA or mock) and harvested after 24 h. For relative expression levels the ΔCt values per gene/sample were determined relative to *AT1G13320*. The plot shows -ΔCt values that are normalized so that the mean -ΔCt value of mock-treated Col-0 is 0. Asterisks indicate significant differences of the mutants with Col-0 within a treatment (ANOVA followed by Dunnett’s test; *n*=5 (a) or 3 (b). * *P* < 0.05, ** *P* < 0.01, *** *P* < 0.001). Points that lie beyond 1.5 times the interquartile range are shown as dots in the plot, and were taken along with the other points in the statistical analysis. See Fig. S1 (File S1) for expression of *PR1* and *VSP2*. Experiment was repeated with similar results.

### Two cysteine mutations in NPR1 specifically affect SA/JA crosstalk

Cysteine-based modifications of NPR1 are critical for its function (Spoel and Loake 2011; Kumar et al. 2022). In the transgenic lines *C82A* and *C216A (*in the *npr1-1* background), mutations of Cys^82^ or Cys^216^ to alanine result in reduced oligomerization and increased nuclear localization of NPR1 (Mou et al. 2003; Kumar et al. 2022). Based on our results presented above, we hypothesized that this increased nuclear localization of NPR1 would not reduce and may even increase repression of JA-responsive transcription by SA. To address this, we performed experiments on mature plants using the same hormone dipping setup as our nuclear localization experiments. We found that *PR1* induction at 5 h and 24 h after SA + MeJA treatment was not significantly altered in either of the NPR1 cysteine mutants compared to Col-0 (Fig. 2a; File S1 - Fig. S2a). This suggests that some core functions of the SA pathway were unaffected by these mutations. However, *PDF1.2* expression after SA + MeJA significantly differed between Col-0 and the two cysteine mutants: while in Col-0 it was lower compared to the elevated level that was induced by MeJA treatment alone, in the two cysteine mutants it was similar or even slightly higher in the double treatment compared to the single treatment (Fig. 2b). A similar trend was observed for *VSP2* expression (File S1 - Fig. S2b). These results show that while expression of *NPR1^C82A^*and *NPR1^C216A^* can mostly complement the *PR1* activation function of NPR1 in the *npr1-1* mutant background, it does not restore SA-mediated suppression of JA-responsive genes. This suggests that Cys^82^ and Cys^216^ are specifically required for NPR1’s role in SA/JA crosstalk.

**Fig. 2.**
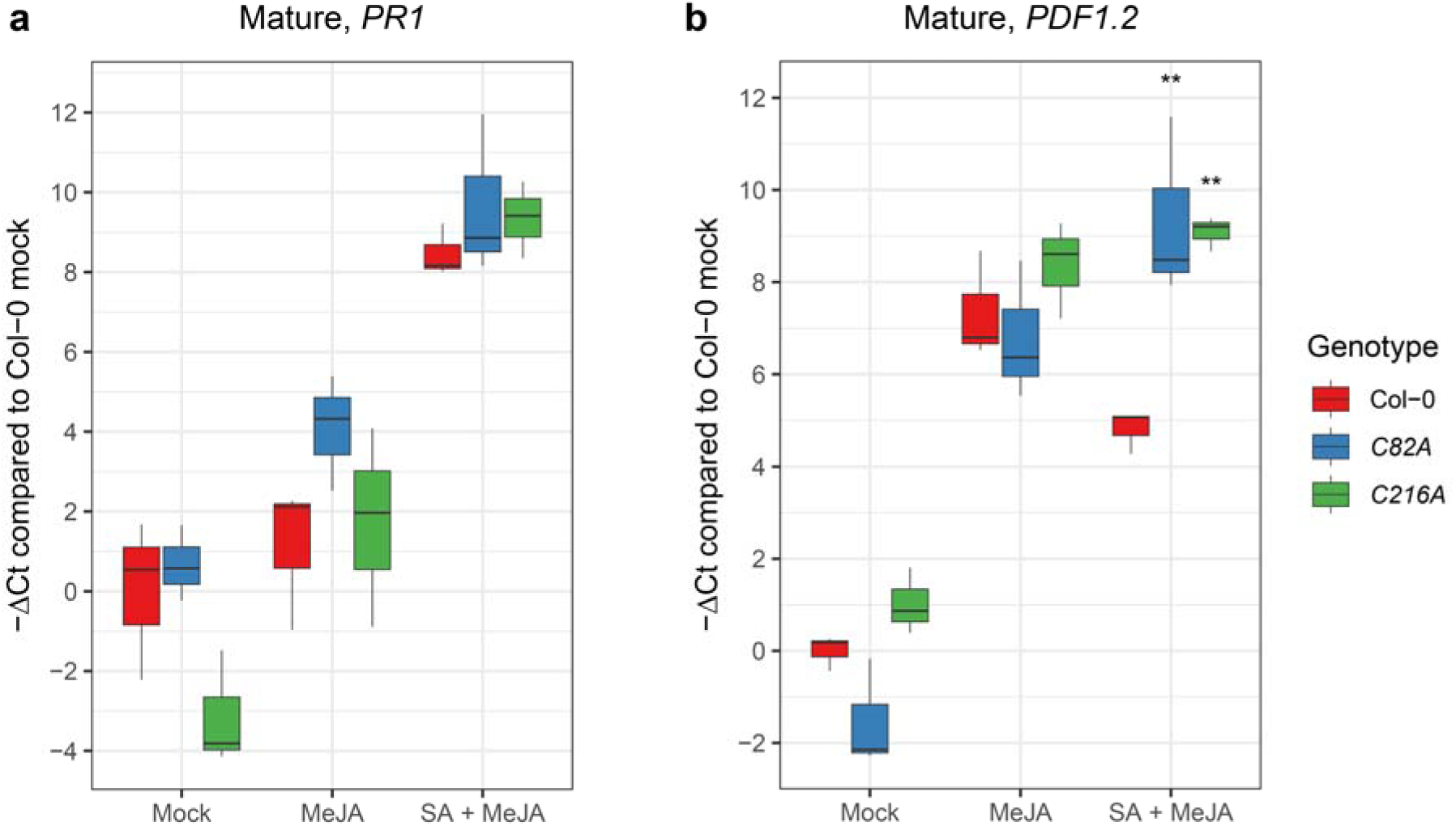
Effects of mutations in NPR1 cysteine residues on SA/JA crosstalk. qRT-PCR expression analysis of (**a**) SA marker gene *PR1* and (**b**) JA marker gene *PDF1.2* after hormone treatments in various NPR1 cysteine mutants. Gene expression was measured in 5-week-old plants in pots. Rosettes were dipped in hormone solution (0.1 mM MeJA and/or 1 mM SA or mock) and harvested after 24 h. For relative expression levels the ΔCt values per gene/sample were determined relative to *AT1G13320*. The plot shows -ΔCt values that are normalized so that the mean -ΔCt value of mock-treated Col-0 is 0. Asterisks indicate significant differences of the mutants with Col-0 within a treatment (ANOVA followed by Dunnett’s test; *n*=3. ** *P* < 0.01). See Fig. S2 (File S1) for expression of *PR1* and *VSP2* at 5 h after treatment. Experiment was repeated with similar results.

### RNA-seq of NPR1 mutants reveals WRKY TFs as potential regulators of SA/JA crosstalk

To investigate how nuclear NPR1 mediates suppression of JA responses, we examined processes affected by the *C82A* and *C216A* NPR1 mutants that may underlie the observed impairment in SA/JA crosstalk. We hypothesized that the cysteine mutations in NPR1 alter its properties such that it can no longer interact with specific TFs that are directly or indirectly required for repressing JA signaling, while (possibly) altered interactions with other SA regulators have smaller effects on SA signaling. The cysteine mutants would therefore be more affected in expression of crosstalk-related genes than of other SA-related genes. To test this, we performed RNA-seq analysis on Col-0 under basal conditions and on Col-0, *npr1-1*, and mutants *C82A* and *C216A* after SA + MeJA treatment, creating conditions where SA/JA crosstalk is active in Col-0 but not in the mutants. For each of the three biological replicates, developmental leaf number 8 of one plant was harvested at 5 h after treatment. First, we verified expression of the different *NPR1* alleles. As shown in Fig. S3a (File S1), the expression of wild-type *NPR1* was 2.5 times upregulated in Col-0 after SA + MeJA treatment, but undetectable in *npr1-1*, *C82A*, and *C216A*. We also confirmed the presence of the *npr1-1* mutation in the *npr1-1* and the *C82A* and *C216A* mutants. In all biological replicates of the *C82A* and *C216A* lines, in which C>A mutant versions of *NPR1* are overexpressed in the *npr1-1* background, the C>A variants of *NPR1* were indeed higher expressed compared to the wild-type *NPR1* version in Col-0 (File S1 - Fig. S3a). However, we noticed that the different replicates (i.e., individual plants) of *C82A* and *C216A* expressed the C>A mutant versions to different extents. In two replicates of *C82A* and one of *C216A*, the C>A *NPR1* transcript levels were around 15-to 25-fold overexpressed, whereas in the other replicates the induction was around 5-fold (File S1 - Fig. S3a).

We next examined whether the variation in C>A *NPR1* transcript abundance was associated with differences in the overall transcriptional response, using principal component analysis (PCA) of the RNA-seq profiles from all samples, including all C82A and C216A biological replicates. We found that indeed, the transcriptome pattern in the replicates with relatively high C>A *NPR1* transcript level (*C82A-2*, *C82A-3*, and *C216A-3*) resembled that of SA + MeJA-treated Col-0 (File S1 - Fig. S4), including SA-induced elevated expression of *PR1* (File S1 - Fig. S3B), whereas the replicates with lower C>A *NPR1* expression (*C82A-1*, *C216A-1*, and *C216A-2*) resembled mock-treated Col-0 and SA + MeJA-treated *npr1-1* (File S1 - Fig. S4), including no induction of *PR1* expression (File S1 - Fig. S3). We thus speculated that expression of C>A-mutated *NPR1* in samples *C82A-1*, *C216A-1*, and *C216-2* was too low to complement SA signaling in the *npr1-1* mutant. Therefore, the transcript profiles of these replicates were excluded from follow-up analyses.

We analyzed in closer detail the two replicates of the *C82A* line that showed high expression of the C>A variant of NPR1. We compared the differentially expressed genes in SA + MeJA-treated Col-0 plants relative to SA + MeJA-treated *npr1-1* or replicate 2 and 3 of SA + MeJA-treated *C82A* (*C82A-2/3*). We first investigated which section of the NPR1-affected transcriptome was also affected in *C82A-2/3*. In SA + MeJA-treated *npr1-1*, 5499 genes were significantly differentially expressed compared to SA + MeJA-treated Col-0, divided into 3019 higher and 2480 lower expressed genes (*P* ≤ 0.05; Data Set S1). This large number underlines the major role of NPR1 in the SA gene regulatory network and SA/JA crosstalk. In contrast, in SA + MeJA-treated *C82A-2/3* plants, only 97 genes were significantly differentially expressed compared to SA + MeJA-treated Col-0 (*P* ≤ 0.05; Data Set S1). This shows that only a small part of the transcriptome is affected by the mutation in Cys^82^. Of these genes, 46 were lower expressed in *C82A-2/3* than in Col-0, of which 40 were also lower expressed in SA + MeJA-treated *npr1-1* compared to SA + MeJA-treated Col-0. This suggests that the C>A mutation in Cys^82^ of NPR1 results in reduced expression of a small subset of genes whose expression is dependent on NPR1, but does not affect the majority of NPR1-dependent genes.

Genes that are upregulated by SA + MeJA in Col-0 but significantly less so in both *C82A-2/3* and *npr1-1* are potential candidates to play a role in NPR1-regulated SA/JA crosstalk. In Col-0, SA + MeJA treatment induced the expression of 33 of the 46 genes that were lower expressed in SA + MeJA-treated *C82A-2/3* than in SA + MeJA-treated Col-0. The expression of 32 of these genes (excluding *ARACNIN1*) was also significantly lower in *npr1-1* compared to Col-0, confirming their NPR1 dependency (Fig. 3). For many of these genes the function is still unknown. Interestingly, four genes affected in *C82A-2/3* encode TFs that have been described to be direct transcriptional targets of NPR1 (Wang et al. 2006). These are *WRKY18*, *WRKY38*, *WRKY53*, and *WRKY70* (indicated with a red dot in Fig. 3). This suggests that the *C82A* mutation in NPR1 leads to reduced transcriptional activation of these four genes by NPR1.

**Fig. 3.**
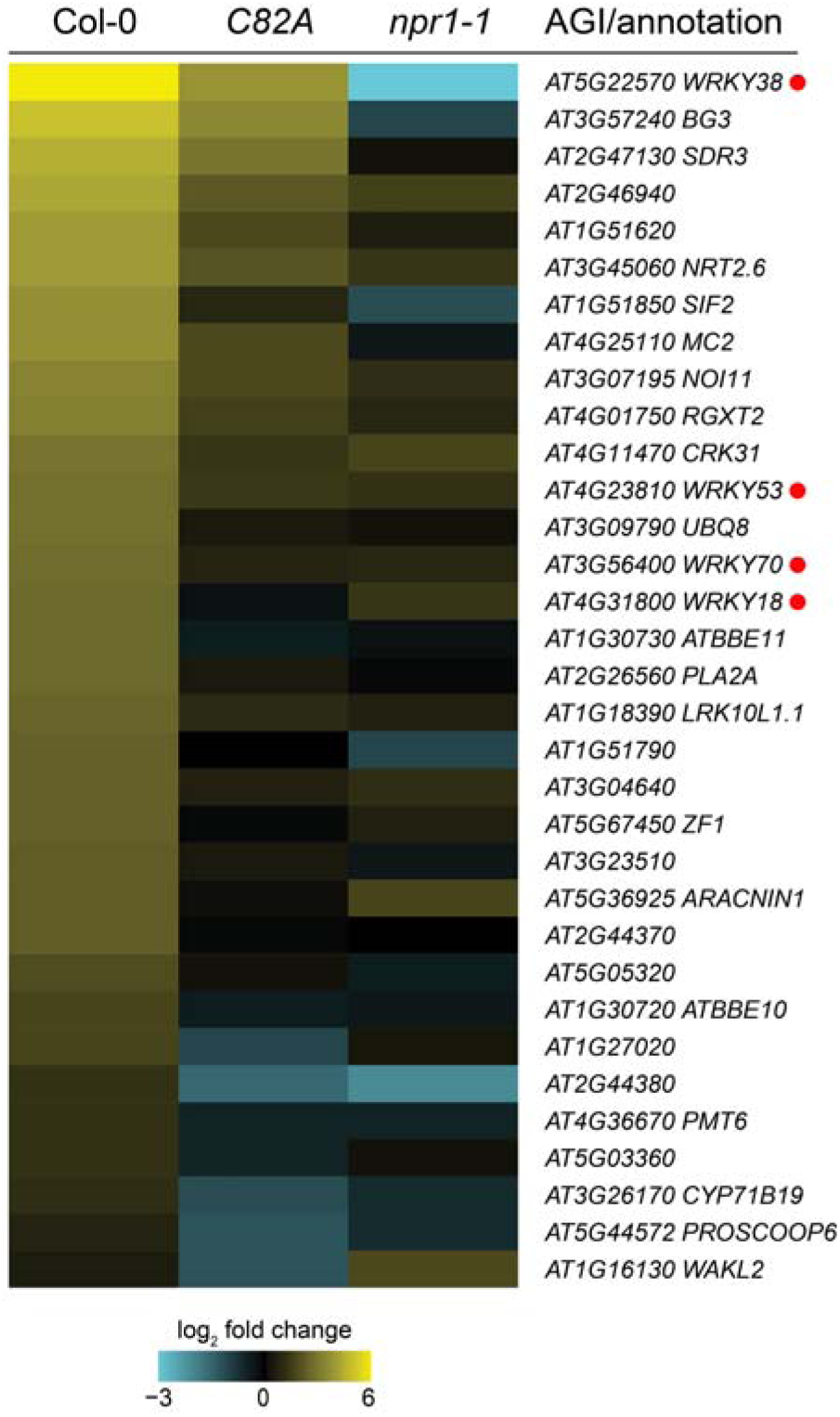
Reduced SA + MeJA-induced expression of genes in NPR1 cysteine mutant *C82A.* Heat map representation of gene expression in Col-0 (*n=3*), *C82A-2/3* (*n*=2) and *npr1-1* (*n*=3) plants measured with RNA-seq. Shown is the log_2_-fold change in expression of genes in SA + MeJA-treated plants (Col-0, *C82A-2/3* or *npr1-1*) relative to mock-treated Col-0 plants. The 33 genes depicted here were significantly lower expressed in *C82A-2/3* compared to Col-0 (adjusted *P* ≤ 0.05) and induced by SA + MeJA in Col-0 (log_2_-fold change (SA + MeJA)/mock Col-0 ≥ 0.6). Genes are identified with AGI and (where available) TAIR annotation. A red dot indicates that the gene is reported as a direct NPR1 target (Wang et al. 2006). See Data Set S1 for all differentially expressed genes between *npr1-1* and Col-0 and between *C82A-2/3* and Col-0.

### WRKY53 represses PDF1.2 expression downstream of ORA59 transcription

Reduced expression of *WRKY18, WRKY38, WRKY53* and *WRKY70* may explain the lack of SA/JA crosstalk in the *C82A* mutant, as WRKY53 and WRKY70 have previously been implicated in regulating this interaction (Hu et al. 2012; Jiao et al. 2022; Li et al. 2006). We designed a transient protoplast assay to examine whether any of these WRKY TFs can suppress JA signaling. We chose to focus on the effect of these WRKY TFs on the ERF branch (measured by the marker gene *PDF1.2*), since the effect of NPR1 on the MYC branch was already found to be dependent on the physical NPR1-MYC2 interaction (Nomoto et al. 2021). We constructed a single plasmid expressing multiple genes for this assay (Fig. 4a). It contained the *PDF1.2* promoter fused to the β*-glucuronidase* (*GUS*) gene as a readout of ERF branch activity, and *ORA59* under a constitutive 35S promoter to drive ERF branch activation. This setup allowed us to examine crosstalk mechanisms downstream of *ORA59* transcription, the level at which SA suppresses *PDF1.2* in whole plants (He et al. 2017; Van der Does et al. 2013). We also included the luciferase gene from firefly (*LucF*) expressed from the 35S promoter to normalize the GUS signal for the number of transfected protoplasts. Additionally, the construct contained a candidate WRKY TF expressed from the 35S promoter, or an empty module as a control. We also added *Enhanced Yellow Fluorescent Protein* (*EYFP*) expressed from the 35S promoter to the construct to verify transfection of the protoplasts during our experimental procedure. As expected, *ORA59* overexpression strongly induced *PDF1.2* expression compared to the control where an empty module was put in the place of *35S_pro_:ORA59* (Fig. 4b). Notably, overexpression of *WRKY53* coexpressed with *ORA59* significantly reduced *PDF1.2* expression compared to *ORA59* alone. Overexpression of *WRKY70* also significantly reduced *PDF1.2* expression, though less consistently over independent experiments (Fig. 4b; File S1 - Fig. S5) and to a lesser extent than *WRKY53*, which consistently and strongly suppressed *PDF1.2* in three independent experiments (Fig. 4b; File S1 - Fig. S5). Overexpression of *WRKY18* or *WRKY38* showed no meaningful effect on *PDF1.2* (Fig. 4b; File S1 - Fig. S5), with only a minor suppression by *WRKY18* in one experiment (File S1 - Fig. S5a). Altogether, our results suggest that WRKY53, and possibly WRKY70, is involved in SA/JA crosstalk by downregulating JA-responsive gene expression downstream of *ORA59* transcription.

**Fig. 4.**
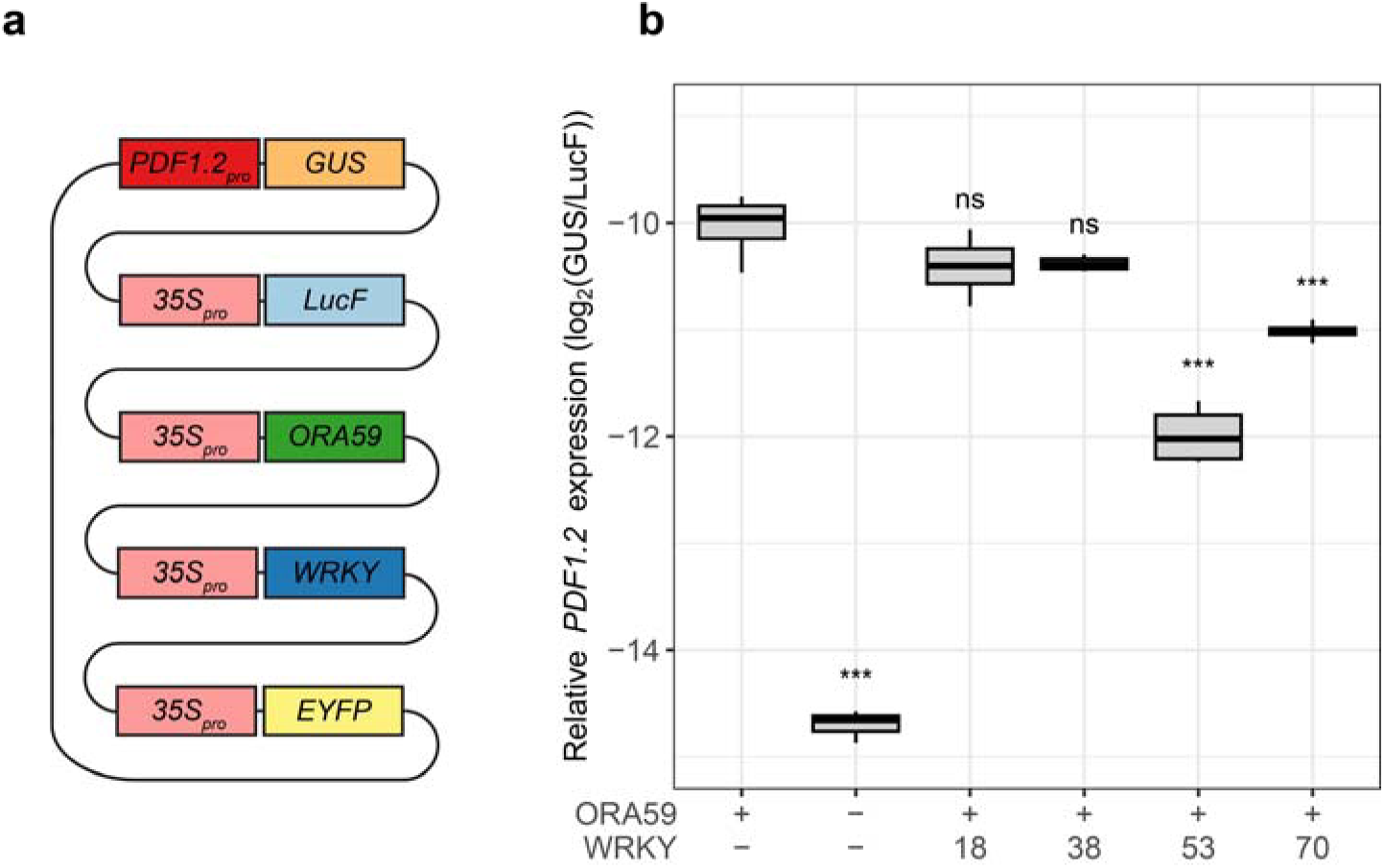
Suppression of ORA59-induced *PDF1.2* expression by WRKY TFs in Arabidopsis protoplasts. (**a**) Schematic representation of the plasmids used in the experimental setup. Protoplasts isolated from 3-week-old plants were transfected with a single plasmid containing the *GUS* gene expressed from the *PDF1.2* promoter, the *LucF* gene expressed from the 35S promoter and the *EYFP* gene expressed from the 35S promoter. Additionally, constructs contained either the *ORA59* gene expressed from the 35S promoter (‘ORA59 +’ in panel b) or only the 35S promoter and terminator (‘ORA59 –‘ in panel b), and they contained one of four WRKY TFs (‘WRKY 18, 38, 53 or 70’ in panel b) expressed from the 35S promoter, or only the 35S promoter and terminator (‘WRKY –’ in panel b). Protoplasts were harvested approximately 20 h after transfection. (**b**) Relative *PDF1.2* expression was determined as the log_2_ of relative GUS levels divided by relative LucF levels, as quantified using the substrates MUG and luciferin, respectively. Asterisks indicate significant differences from the reference (ORA59+ WRKY-) according to Dunnett’s multiple-comparison test (\**P* < 0.05, \*\**P* < 0.01, \*\*\**P* < 0.001, ns = not significant; *n*=4). See Fig. S5 (File S1) for two other repeats of the same experiment.

### Four additional WRKY TFs suppress ORA59-induced PDF1.2 expression

Regulation of plant processes is often carried out by redundantly functioning proteins. For example, WRKY53 is known to operate redundantly with WRKY46 and WRKY70 in basal resistance against *Pseudomonas syringae* and likely also in suppression of *PDF1.2* (Hu et al. 2012). We therefore considered the possibility that other WRKY TFs related to WRKY53 are also involved in suppression of ORA59-induced *PDF1.2* expression. WRKY53 is part of a clade of WRKY TFs known as group III (Kalde et al. 2003; File S1 - Fig. S6). The majority of TFs in this group are induced by SA (Hickman et al. 2019) and a number of TFs were already implicated in SA/JA crosstalk (Caarls et al. 2015; Kalde et al. 2003). We therefore decided to analyze all TFs in this clade that were induced by SA according to Hickman et al. (2019), as well as other WRKY TFs known to suppress JA signaling according to Caarls et al. (2015). We tested this in the same protoplast system as before so that we could assess which of these WRKY TFs suppress *PDF1.2* expression downstream of *ORA59* transcription. We did not include WRKY55 and WRKY64 in this experiment because we were unable to clone these two genes. In two independent runs of this experiment, ORA59 again strongly induced *PDF1.2* expression and WRKY53 suppressed it, as expected (Fig. 5; File S1 - Fig. S7). WRKY70 also suppressed *PDF1.2* expression, although only significantly in one of the two experimental runs (Fig. 5) and to a smaller extend than WRKY53. Four other WRKY TFs were able to suppress ORA59-induced *PDF1.2* expression in both experimental runs: WRKY50, WRKY54, WRKY63 and most notably WRKY67 (Fig. 5; File S1 - Fig. S7). Of these TFs, WRKY50 (Gao et al. 2011) and WRKY54 (Li et al. 2017) were previously associated with suppression of JA signaling, but to the best of our knowledge WRKY63 and WRKY67 were not. In our RNA-seq data set of SA + MeJA-treated Col-0 and *npr1-1* plants (Data Set S1), *WRKY50*, *WRKY53* and *WRKY54* were differentially expressed between Col-0 and *npr1-1*, suggesting they act in NPR1-dependent SA/JA crosstalk. Conversely, *WRKY63* and *WRKY67* were not differentially expressed, suggesting that their role in SA/JA crosstalk may be independent of NPR1. There was no clear phylogenetic relationship between the suppressing WRKY TFs within group III of the WRKY TF family (File S1 - Fig. S6).

**Fig. 5.**
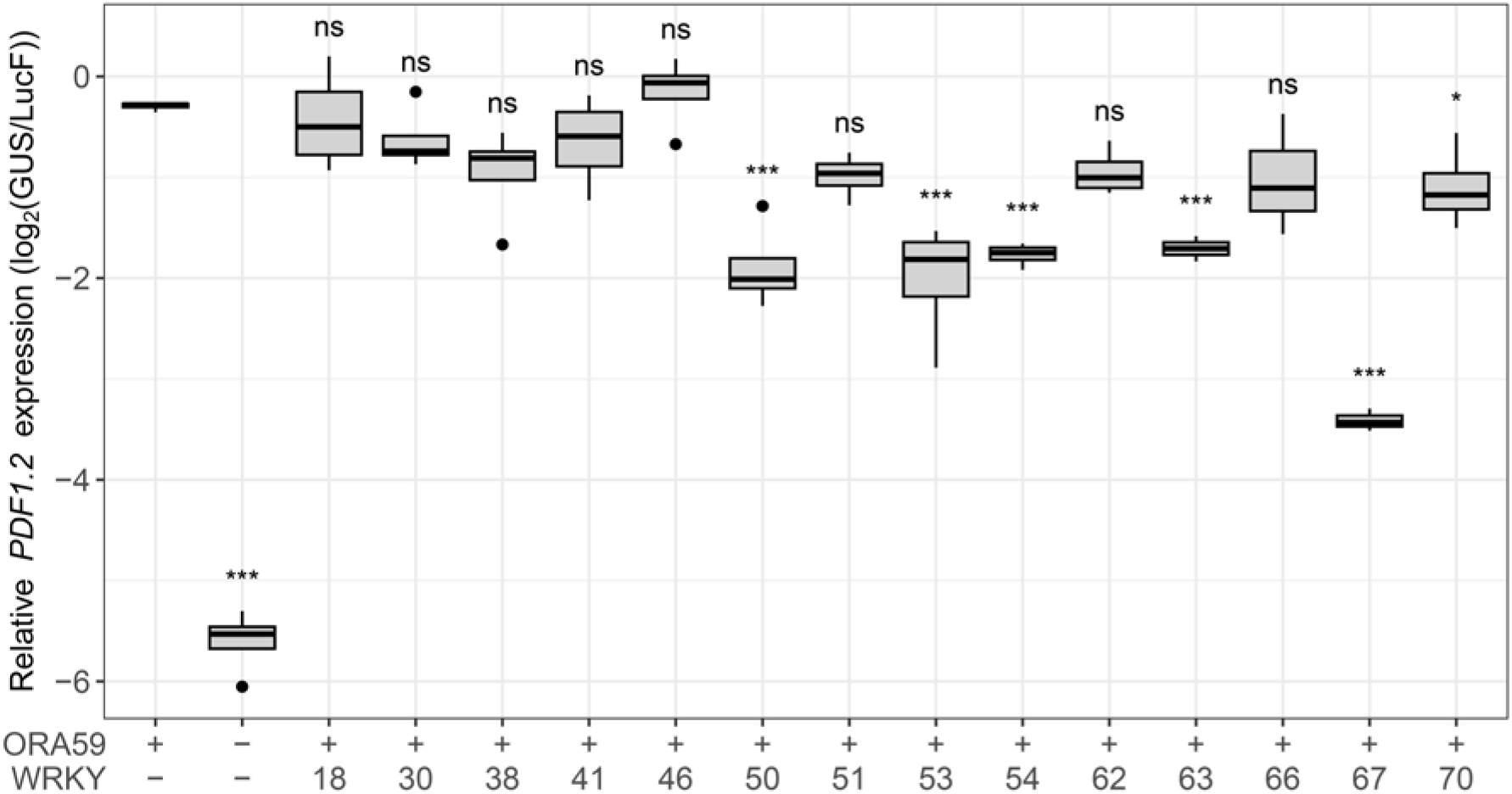
Suppression of ORA59-induced *PDF1.2* expression by selected WRKY TFs in Arabidopsis protoplasts. Protoplasts isolated from 3-week-old plants were transfected with a single plasmid, depicted and described in (the legend of) Fig. 4a. Relative *PDF1.2* expression was determined as the log_2_ of relative GUS levels divided by relative LucF levels, as quantified using the substrates MUG and luciferin, respectively. Asterisks indicate significant differences from the reference (ORA59+ WRKY-) according to Dunnett’s multiple-comparison test (\**P* < 0.05, \*\**P* < 0.01, \*\*\**P* < 0.001, ns = not significant; *n*=4). Points that lie beyond 1.5 times the interquartile range are shown as dots in the plot, and were taken along with the other points in the statistical analysis. A repeat of this experiment is depicted in Fig. S7 (File S1).

### WRKY53 and WRKY67 regulate SA/JA crosstalk by reducing ORA59 protein levels

SA is known to suppress ORA59-induced *PDF1.2* expression by inducing degradation of ORA59 protein (Van der Does et al. 2013; He et al. 2017). We hypothesized that the WRKY TFs we identified as suppressors may (indirectly) affect ORA59 protein stability. To test this, we used our protoplast system, but linked *ORA59* to the *Renilla reniformis* luciferase gene (*LucR*) to allow the ratio LucR to LucF to serve as a proxy for ORA59 protein levels (Fig. 6a). Due to the large size of the resulting plasmid, the WRKY TFs were expressed from a separate plasmid, and the protoplasts were cotransfected with both plasmids (Fig. 6a). Consistent with our previous findings (Fig. 5), overexpression of all five WRKY TF genes (*WRKY50*, *WRKY53*, *WRKY54*, *WRKY63*, and *WRKY67*) reduced *PDF1.2* expression, validating the effectiveness of dual-plasmid transfection (Fig. 6b). Importantly, overexpression of *WRKY53* and *WRKY67* led to a small but significant decrease in ORA59 protein levels (Fig. 6), despite its expression from a constitutive promoter. This effect of WRKY53 and WRKY67 was reproducible in independent experiments, although in one of three independent experiments the reduction of ORA59 protein levels upon overexpression of WRKY53 was not significant (File S1 - Fig. S8). Together, our results suggest that WRKY53 and WRKY67 contribute to suppression of *PDF1.2* expression partly by targeting ORA59 protein levels.

**Fig. 6.**
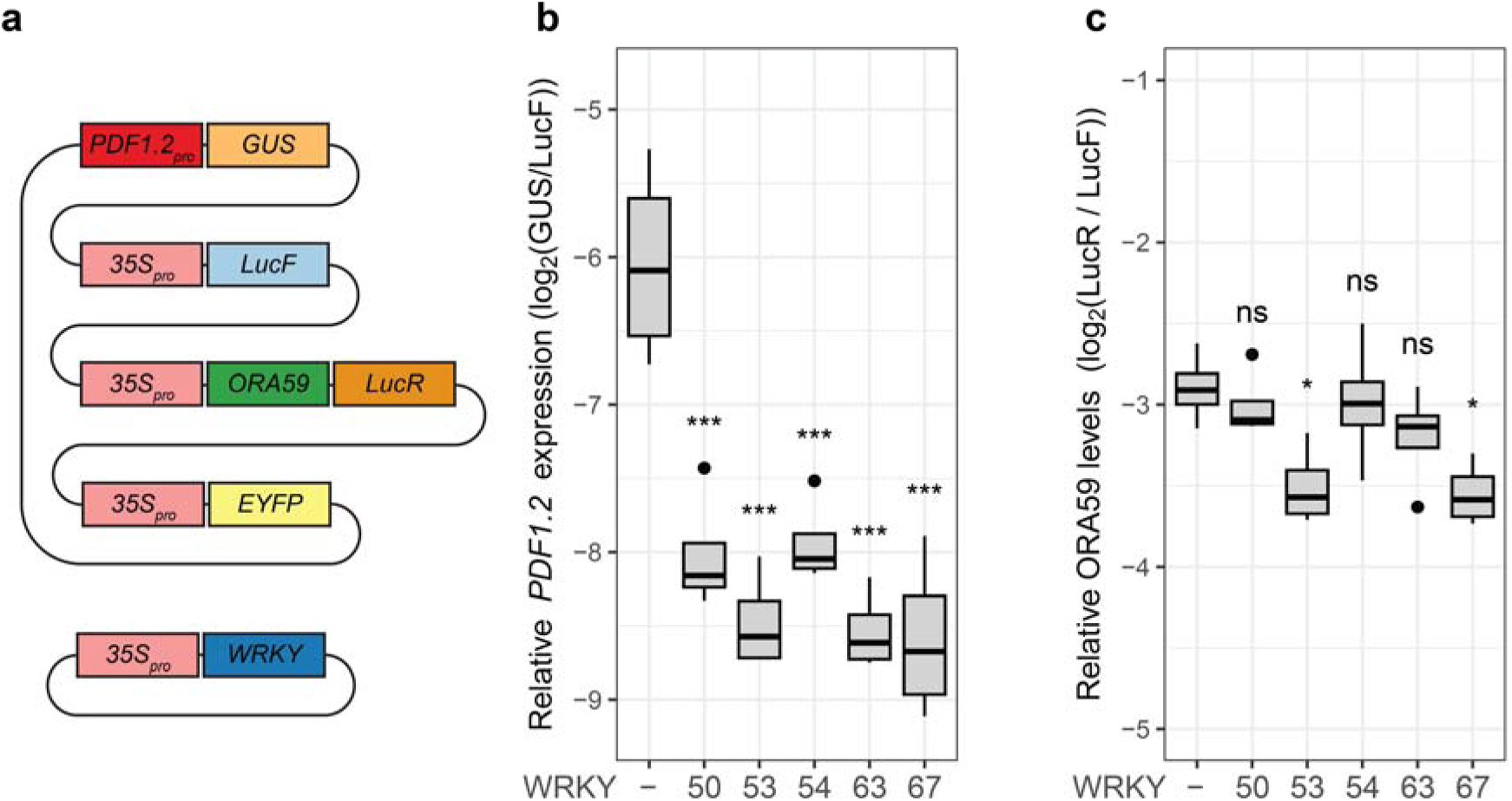
Suppression of *PDF1.2* expression and relative ORA59 protein levels by selected WRKY TFs. (**a**) Protoplasts isolated from 3-week-old plants were transfected with two plasmids. The first contained the *GUS* gene expressed from the *PDF1.2* promoter, the *LucF* gene expressed from the 35S promoter, the *ORA59* gene fused to the *LucR* gene expressed from the 35S promoter, and the *EYFP* gene expressed from the 35S promoter. The second plasmid contained a WRKY TF (indicated with its number in panels b and c) expressed from the 35S promoter, or only the 35S promoter and terminator (‘WRKY –‘ in panels b and c). Protoplasts were harvested approximately 20 h after transfection. (**b**) Relative *PDF1.2* expression was determined as the log_2_ of relative GUS levels divided by relative LucF levels, as quantified using the substrates MUG and luciferin, respectively. (**c**) Relative ORA59 levels were determined as the log_2_ of relative LucR levels divided by relative LucF levels, as quantified using the substrates coelenterazine and luciferin, respectively. Asterisks in (**b**) and (**c**) indicate significant differences from the reference (‘WRKY-‘) according to Dunnett’s multiple-comparison test (\**P* < 0.05, \*\**P* < 0.01, \*\*\**P* < 0.001, ns = not significant; *n*=4).). Points that lie beyond 1.5 times the interquartile range are shown as dots in the plot, and were taken along with the other points in the statistical analysis. Repeats of this experiment with only WRKY53 or only WRKY67 are shown in Fig. S8 (File S1).

## DISCUSSION

NPR1 is an important regulator of SA-induced gene expression and of SA/JA crosstalk. While regulation of NPR1 in response to SA has been studied extensively, NPR1’s role in SA/JA crosstalk, particularly on the ERF branch of the JA pathway, remains less understood. This study investigated the function of NPR1 in SA/JA crosstalk, focusing on its nucleo-cytoplasmic localization and downstream transcriptional effects. In our experimental setups, cytosolic NPR1 did not suppress the JA marker gene *PDF1.2* following SA treatment in both seedlings and mature plants (Fig. 1) and hence, nuclear localization of NPR1 appeared essential for SA/JA crosstalk. As this contradicts findings by Spoel et al. (2003), the role of nuclear/cytosolic NPR1 in regulating SA/JA crosstalk is likely context-dependent, modulated by factors that are currently unknown. Additionally, we tested two lines overexpressing cysteine-to-alanine-mutated versions of NPR1, *C82A* and *C216A*, which turned out to be impaired in SA-mediated suppression of *PDF1.2*, but to display normal levels of SA-induced *PR1* expression. Our RNA-seq analysis of SA + MeJA treatment revealed that only 97 of 5499 NPR1-dependent genes were affected in *C82A*. This indicates subtle effects by the cysteine mutation that likely mostly influence genes affected by crosstalk, and that Cys^82^ and Cys^216^ serve as key residues for NPR1-mediated SA/JA crosstalk. We then identified five SA- or SA/NPR1-regulated WRKY TFs that suppressed ORA59-induced *PDF1.2* expression, independent of *ORA59* transcription, in a protoplast system. Of these TFs, WRKY53 and WRKY67 reduced the accumulation of ORA59 protein. Together, this points to a model where SA induces expression of specific crosstalk-regulating WRKY TF genes, for which Cys^82^ and Cys^216^ of NPR1 are important. Other SA-inducible crosstalk-regulating WRKY TFs can be transcriptionally activated in an NPR1-independent pathway, but may still need NPR1 for their crosstalk effect. The encoded WRKY TFs can suppress ORA59-activated *PDF1.2* expression partly via reducing accumulation of ORA59 (Fig. 7).

**Fig. 7.**
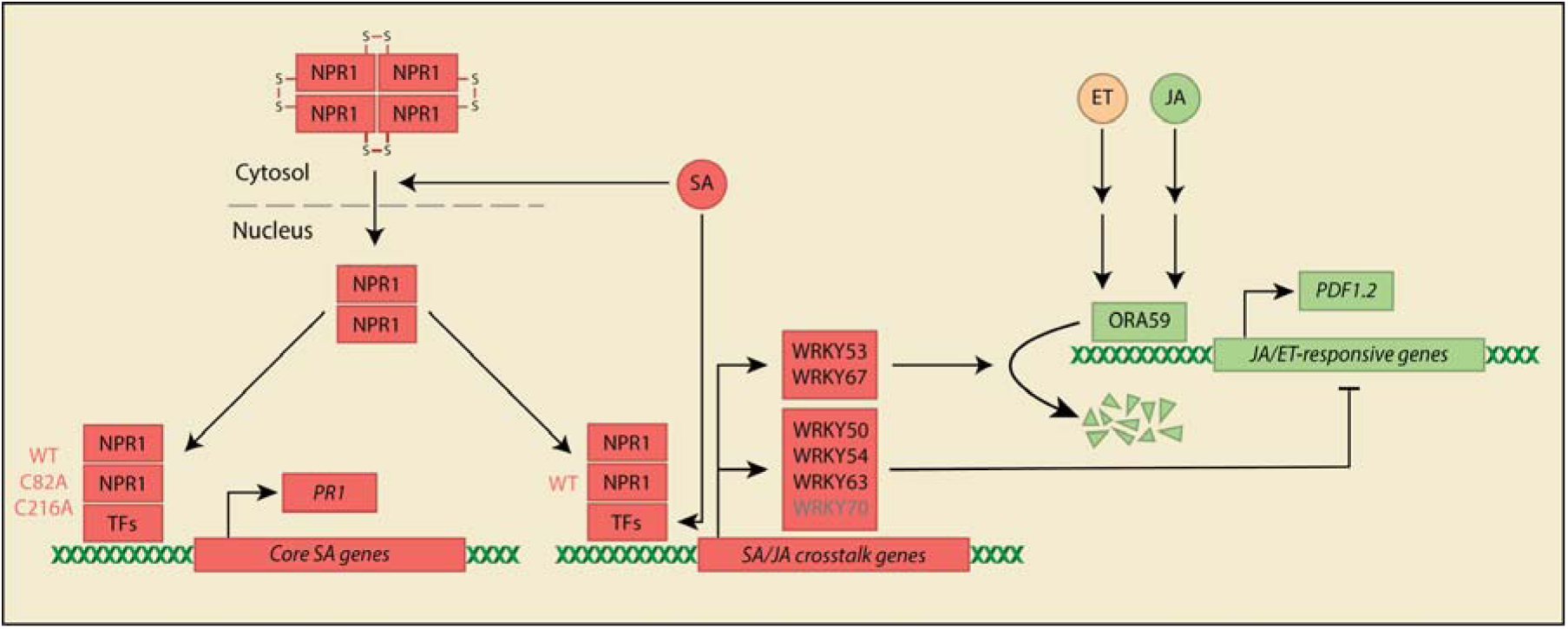
Schematic model of the role of NPR1 and various WRKY proteins in repression of the ERF branch of the JA pathway. NPR1 resides in the cytosol as an oligomer held together by disulfide bridges between cysteine residues such as Cys^82^ and Cys^216^. SA triggers a redox change, which causes reduction of these cysteine residues, leading to decreased oligomerization and increased nuclear localization of NPR1. In the nucleus, NPR1 forms a homodimer and acts as a transcriptional cofactor. Together with specific TFs it activates core SA genes, such as *PR1*, and SA/JA crosstalk regulatory genes, including specific WRKY TF genes. In the *C82A* and *C216A* mutants NPR1 is only able to activate core SA genes such as *PR1*, but not crosstalk-specific WRKY TF genes. SA can also induce expression of specific WRKY TF genes independently of NPR1. When ET and JA levels are high, they activate *ORA59* together, which causes upregulation of JA/ET-responsive genes such as *PDF1.2*. WRKY53 and WRKY67 cause reduced accumulation of the ORA59 protein, leading to reduced *PDF1.2* expression. WRKY50, WRKY54, WRKY63 and possibly WRKY70 repress *PDF1.2* through a different, unknown mechanism, acting downstream of *ORA59* transcription.

### Cytosolic NPR1 did not suppress JA marker genes

Previous findings reported suppression of MeJA-induced *PDF1.2* by cytosolic NPR1-HBD following SA treatment (Spoel et al. 2003). Furthermore, the *npr1-3* mutant, which lacks the C-terminal part of the NPR1 protein that contains the nuclear localization signal, exhibited reduced but notable SA-mediated suppression of *PDF1.2* (Leon-Reyes et al. 2009). These results suggested a cytoplasmic role of NPR1 in SA/JA crosstalk. However, under the experimental conditions of the current study, two genotypes expressing cytosolic NPR1 (*NPR1-HBD* and *npr1-nls*) were impaired in SA-mediated suppression of *PDF1.2* and – to a lesser extent – *VSP2* in both adult plants and seedlings (Fig. 1; File S1 - Fig. S1). These results indicate that, in this study, NPR1’s function in SA/JA crosstalk is mostly nuclear, suggesting that the cytoplasmic function of NPR1 in SA/JA crosstalk is context dependent. A nuclear function of NPR1 in SA/JA crosstalk was also found by Nomoto et al. (2021), who found in seedlings that NPR1 directly represses MYC2 by disrupting its binding with MED25. The fact that the nuclear mechanism elucidated in our study does not involve MYC2 underlines that nuclear-located NPR1 can regulate crosstalk via multiple mechanisms.

### Cysteine-to-alanine mutations in NPR1 disrupt SA-mediated suppression of JA marker genes

Mutating cysteine residues Cys^82^ or Cys^216^ into alanine in NPR1 caused the transgenic *C82A* and *C216A* lines that overexpress these mutated versions of the NPR1 protein in the *npr1-1* background to lose suppression of MeJA-induced *PDF1.2* and *VSP2* by SA (Fig. 2b; File S1 - Fig. S2b). Interestingly, similar mutations in rice NPR1 (Cys^76^ and Cys^216^ changed to alanine) have also been shown to prevent suppression of JA-dependent genes and defense (Yuan et al. 2007), supporting the relevance of these cysteine residues for NPR1’s role in SA/JA crosstalk. As the Cys-mutated NPR1 in rice was constitutively localized to the nucleus, these results were interpreted as evidence for a cytosolic role of NPR1 in suppression of JA responses. Mutation of Cys^82^ or Cys^216^ in the Arabidopsis *C82A* and *C216A* lines caused the Cys-mutated NPR1 protein to localize in both the nucleus and the cytosol (Mou et al. 2003). Therefore, the impairment in SA/JA crosstalk in both Cys-mutated NPR1-overexpressing Arabidopsis and rice is unlikely due to exclusion of NPR1 from the cytosol, but rather caused by other effects of the mutations in the Cys residues. For example, the mutations may lead to loss of interaction with other proteins, such as specific TFs, causing reduced activation of particular NPR1 target genes. Unfortunately, we were unable to execute interaction assays with the different NPR1 variants to test this hypothesis.

Kumar et al. (2022) found that NPR1^C82A^, despite being defective in higher-order oligomerization, was still able to homodimerize and could still activate *PR1*, while a fully dimerization-defective mutant was defective in *PR1* induction and resistance to *Pseudomonas syringae* pv. *maculicola* ES4326. Therefore, they concluded that the NPR1 homodimer is the functional unit in activating SA-dependent defense gene expression. Our finding that the *C82A* line was impaired in SA/JA crosstalk suggests that the NPR1^C82A^ homodimer is not functional in regulating suppression of the JA pathway. It is unknown whether this is due to its oligomerization status or to other structural changes that, for example, make the protein unable to interact with certain crosstalk-regulating TFs.

In accordance with Kumar et al. (2022), we found that *PR1* expression in *C82A* was similar to that in Col-0 or even higher, suggesting that core SA signaling was not negatively affected by this mutation (Fig. 2; File S1 - Fig. S3). This is corroborated by our RNA-seq analysis, which showed that only 97 genes were differentially expressed in SA + MeJA-treated *C82A* compared to Col-0, while the *npr1-1* mutation affected 5499 genes. In our study, *WRKY18*, *WRKY38*, *WRKY53*, and *WRKY70* were lower expressed in *C82A* (Fig. 3), potentially linking them to SA/JA crosstalk. Indeed, follow-up experiments implicated WRKY53 and WRKY70 as crosstalk regulators acting downstream of ORA59 transcription in *PDF1.2* suppression. Notably, the C82A-affected WRKY TFs are also known to regulate SA-dependent resistance to biotrophic pathogens and systemic acquired resistance (Spoel et al. 2009; Hu et al. 2012; Powers et al. 2024). This indicates that genes affected by the C82A mutation are not exclusively involved in SA/JA crosstalk, but can also regulate other SA responses. Many of the C82A-affected genes are uncharacterized (Data Set S1) and potentially regulate SA responses, including SA/JA crosstalk, making them interesting candidates for future research.

### At least five WRKY TFs are suppressors of ORA59-induced PDF1.2 expression

Using a protoplast system we found that WRKY53 and possibly WRKY70 are able to repress *PDF1.2* expression downstream of *ORA59* transcription (Fig. 4; File S1 - Fig. S5), confirming earlier reports (Hu et al. 2012; Li et al. 2004). We performed the same protoplast assay with other WRKY TFs of group III (to which WRKY53 and WRKY70 belong) and WRKY TFs that have been reported to suppress JA signaling (Caarls et al. 2015). This indicated that WRKY50, WRKY54, WRKY63 and WRKY67 were also able to suppress ORA59-induced *PDF1.2* expression (Fig. 5). Of the five WRKY TFs that consistently repressed ORA59-induced *PDF1.2* expression, WRKY50, WRKY53 and WRKY54 had previously been linked to suppression of JA signaling, and our RNA-seq data indicated that they likely act in NPR1-dependent crosstalk (Data Set S1). WRKY50 can suppress *PDF1.2* expression together with WRKY51 (Gao et al. 2011), and WRKY53 suppresses *PDF1.2* expression redundantly with WRKY46 and WRKY70 (Hu et al. 2012). Additionally, a recent report suggested that WRKY53 negatively regulates resistance against insects by directly binding to and repressing the promoters of the JA biosynthesis genes *LOX3* and *LOX4* (Jiao et al. 2022). This corroborates that repression of JA signaling is regulated by various mechanisms acting at different levels, further emphasizing the importance of this crosstalk for the plant. As we overexpressed *ORA59* to activate the JA pathway rather than treating with JA, a possible decrease in JA biosynthesis cannot explain our results. Indeed, we found that WRKY53 caused a reduction in ORA59 protein levels (Fig. 6), providing a possible mechanism by which WRKY53 can repress the JA pathway. WRKY54 has also been linked to suppression of the JA pathway, as the *wrky54 wrky70* double mutant displays increased resistance against necrotrophic pathogens and elevated *PDF1.2* expression, as well as higher SA levels (Li et al. 2017). The mechanism hereof was not elucidated. The finding that WRKY63 and WRKY67 can suppress ORA59-induced *PDF1.2* expression is novel, to the best of our knowledge, and our RNA-seq data indicated that they likely act in NPR1-independent crosstalk (Data Set S1). WRKY67 is an especially interesting new candidate regulator of SA/JA crosstalk, as it was found to cause the strongest reduction in ORA59-induced *PDF1.2* expression in protoplasts (Fig. 5) as well as a reduction in ORA59 protein levels (Fig. 6). We found that the group III WRKY TFs that suppressed ORA59-induced *PDF1.2* expression were not phylogenetically related within group III of the WRKY family (File S1 - Fig. S6). This supports the hypothesis that the different WRKY TFs have various modes of action.

Interestingly, we did not observe suppression of ORA59-induced *PDF1.2* expression by WRKY41, WRKY46, WRKY51 and WRKY62, and inconsistent suppression of ORA59-induced *PDF1.2* expression by WRKY70 between experiments, even though all of these WRKY TFs have been implicated in suppression of JA signaling (Caarls et al. 2015). This could indicate that these WRKY TFs act upstream of *ORA59* transcription or require certain cofactors that were not present under our experimental conditions.

### WRKY53 and WRKY67 affect ORA59 protein accumulation

SA is known to repress the ERF branch by destabilizing ORA59 (He et al. 2017; Van der Does et al. 2013). Our protoplast system results indicated that WRKY53 and WRKY67 reduce ORA59 accumulation, potentially by increasing ORA59 protein turnover or by influencing *ORA59* transcript stability or translation efficiency. This raises the prospect of exploring downstream targets of these WRKY TFs, including potential ubiquitin ligases that might influence ORA59 stability. Future research can also include investigation of the targets of the other WRKY TFs to clarify their roles in regulating suppression of ORA59-induced *PDF1.2* expression. Our experimental data suggested that while WRKY53 and WRKY67 reduce ORA59 accumulation, WRKY50, WRKY54 and WRKY63 did not affect it. However, since we do not know if all WRKY TF genes were overexpressed to similar levels and since the effect of WRKY53 and WRKY67 on ORA59 accumulation was small, it is possible that the other tested WRKYs may affect ORA59 accumulation, but that this effect could not be detected in our experimental setup.

In conclusion, this study uncovered a novel nuclear function of NPR1 in mediating SA/JA crosstalk (Fig. 7).. Upon SA-mediated activation, NPR1 translocates to the nucleus, where it promotes the transcription of specific WRKY TF genes involved in SA/JA crosstalk, dependent on residues Cys^82^ and Cys^216^. Some other WRKY TF genes involved in SA/JA crosstalk are also transcriptionally activated by SA, dependently or independently of NPR1 and its Cys82 residue. The crosstalk-regulatory WRKY TFs encoded by these genes subsequently regulate the suppression of JA-responsive genes through various mechanisms, including decreasing the protein levels of the ERF branch regulator ORA59.

## Supporting information

Data Set S1

## ACKNOWLEDGEMENTS

We thank Vince Deosarran for performing pilot experiments with the nuclear localized *npr1* mutant lines and *npr1* cysteine mutant lines. We are grateful for the work of Folkert Sanders, Ellinor Keepers and Raquel Ledo Doval in optimizing our protoplast transfection and GUS and LucF measurement protocols. We are grateful to Xinnian Dong for sharing the *NPR1-HBD*, *npr1-nls, C82A,* and *C216A* seeds.

## AUTHOR INFORMATION

All authors

### Contributions

SCMVW, CMJP, LC, and NA conceived and designed the research. NA, LC, TWV, and BG conducted experiments. RK and BG designed and optimized methodology. MCVV was involved in data analysis. NA, and LC wrote the manuscript, with review and editing from SCMVW and CMJP. All authors read and approved the manuscript.

### Corresponding author

Correspondence to Niels Aerts.

## DATA AVAILABILITY

The sequencing data underlying this article are available in the Short Read Archive (http://www.ncbi.nlm.nih.gov/sra/), and can be accessed with BioProject ID PRJNA1217129.

