## Supplementary material for "Nuclear NPR1 Modulates Salicylic Acid/Jasmonic Acid Crosstalk via Transcriptional Control of WRKY Transcription Factors": File S1

**SUPPLEMENTARY FIGURES**

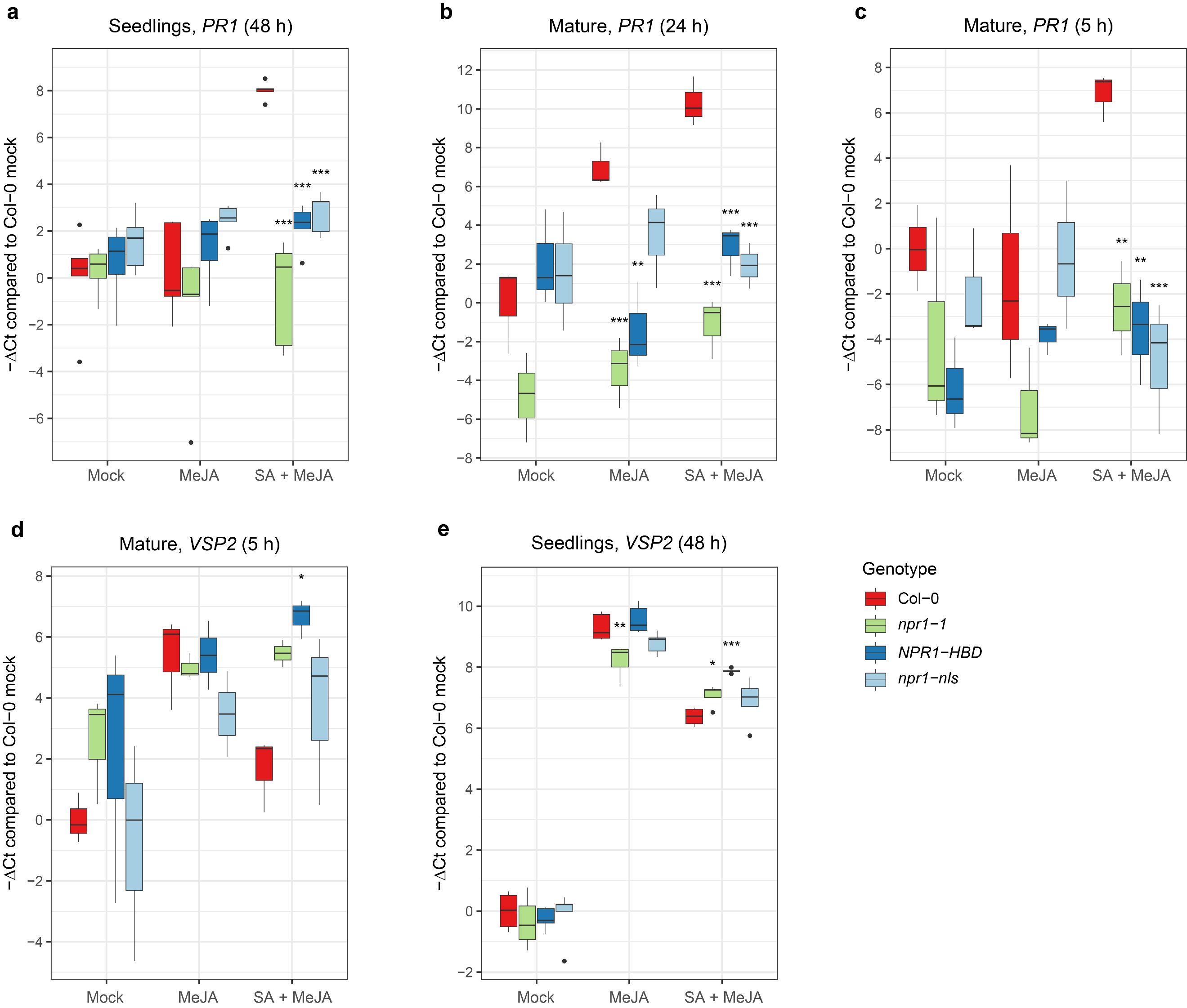

**Fig. S1** Effects of mutations affecting NPR1 localization on SA/JA crosstalk

qRT-PCR expression analysis of (a-c) SA marker gene *PR1* and (d, e) SA marker gene *VSP2* after hormone treatments in various localization-related NPR1 mutants.

(**a, e**) Gene expression measured in 12-day-old seedlings on plates. Plants were treated for 48 h with MeJA (0.02 mM) and SA (0.5 mM).

(**b-d**) Gene expression measured in 5-week-old plants in pots. Rosettes were dipped in hormone solution (0.1 mM MeJA and/or 1 mM SA or mock) and harvested after (d, d) 5 h or (b) 24 h.

For relative expression levels the ΔCt values per gene/sample were determined relative to *AT1G13320*. The plot shows -ΔCt values that are normalized so that the mean -ΔCt value of mock-treated Col-0 is 0 Asterisks indicate significant differences of the mutants with Col-0 within a treatment (ANOVA followed by Dunnett’s test; *n*=5 (for a and e) or *n*=3 (for c-d). * *P* < 0.05, ** *P* < 0.01, *** *P* < 0.001). See Fig. 1 for expression of *PDF1.2*. Points that lie beyond 1.5 times the interquartile range are shown as dots in the plot, and were taken along with the other points in the statistical analysis.

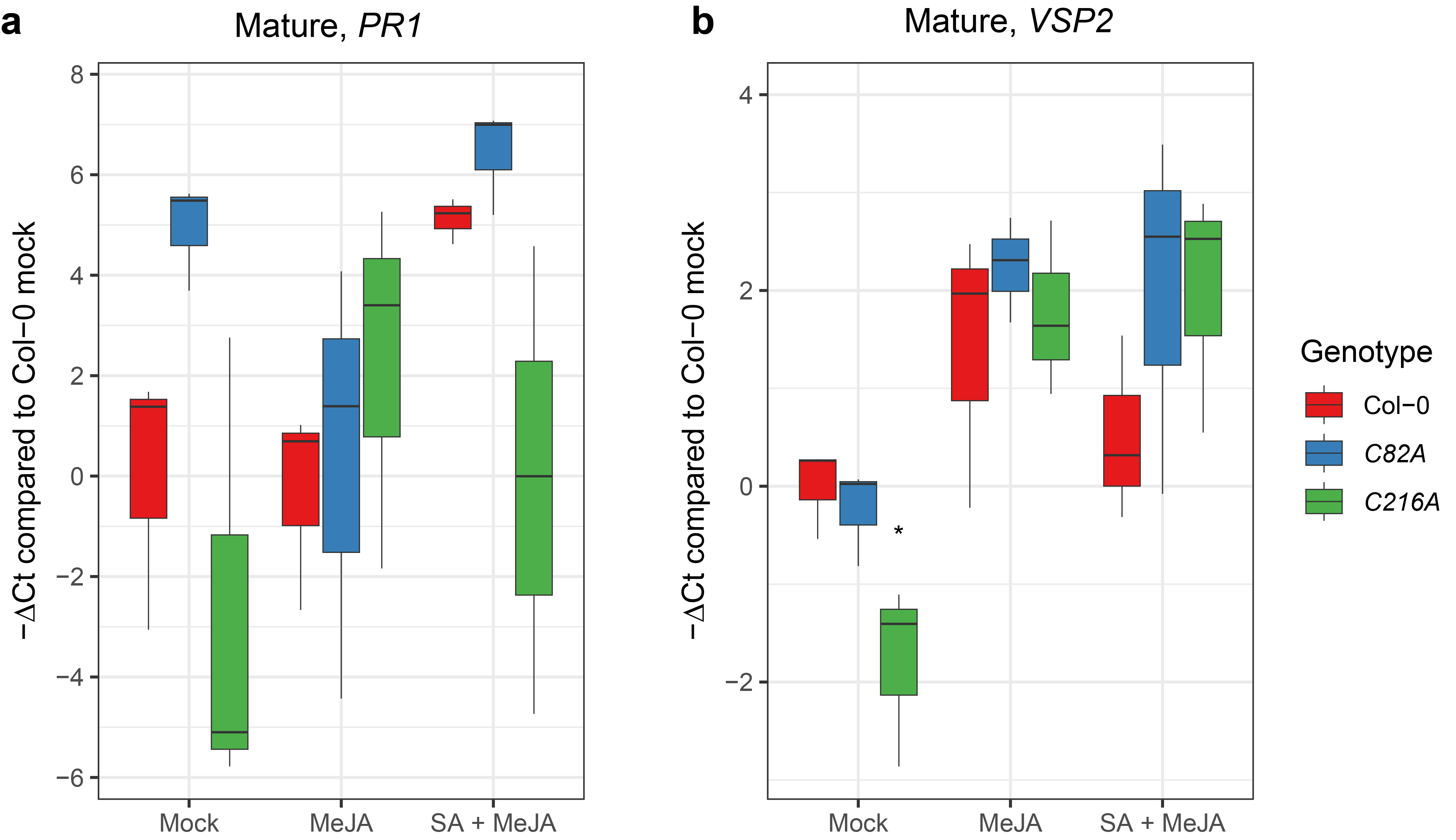

**Fig. S2** Effects of mutations in NPR1 cysteine residues on SA/JA crosstalk

qRT-PCR expression analysis of (**a**) SA marker gene *PR1* and (**b**) JA marker gene *VSP2* after hormone treatments in Col-0 and various NPR1 cysteine mutants. Gene expression was measured in 5-week-old plants in pots. Rosettes were dipped in hormone solution (0.1 mM MeJA and/or 1 mM SA or mock) and harvested after 5 h. For relative expression levels the ΔCt values per gene/sample were determined relative to *AT1G13320*. The plot shows -ΔCt values that are normalized so that the mean -ΔCt value of mock-treated Col-0 is 0. Asterisks indicate significant differences of the mutants with Col-0 within a treatment (ANOVA followed by Dunnett’s test; *n*=5 (a) or 3 (b). * *P* < 0.05). See Fig. 2 for expression of *PDF1.2* and *PR1* at 24 h after treatment.

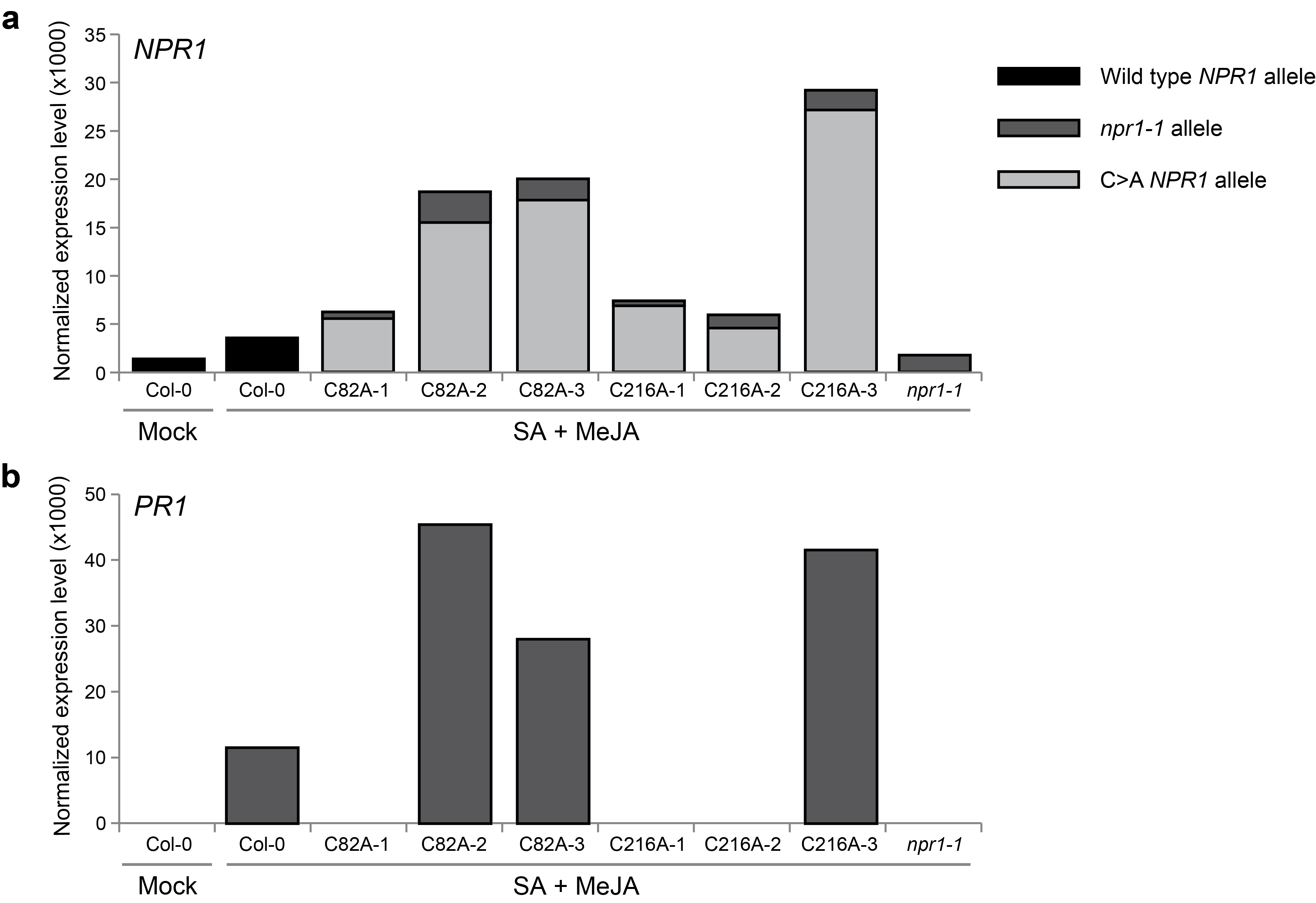

**Fig. S3** Expression levels of *NPR1* alleles and *PR1* in Col-0, *npr1-1* and two NPR1 cysteine mutants revealed by RNA-seq

(**a**) Expression levels of *NPR1* alleles (normalized RNA-seq read counts). Expression of the *npr1-1* allele was estimated by multiplying the total number of *NPR1* counts with the ratio of counts containing the C1000T point mutation of the *npr1-1* allele. The rest of the counts were assigned to the wild-type *NPR1* allele (for Col-0) or the C>A *NPR1* allele (for *C82A* and *C216A*).

(**b**) Expression levels of *PR1* (normalized RNA-seq read counts).

Expression was measured in mock-treated Col-0 plants or SA + MeJA-treated Col-0, *npr1-1* (average of three replicates), and *C82A* and *C216A* mutants (three separate replicates) with RNA-seq.

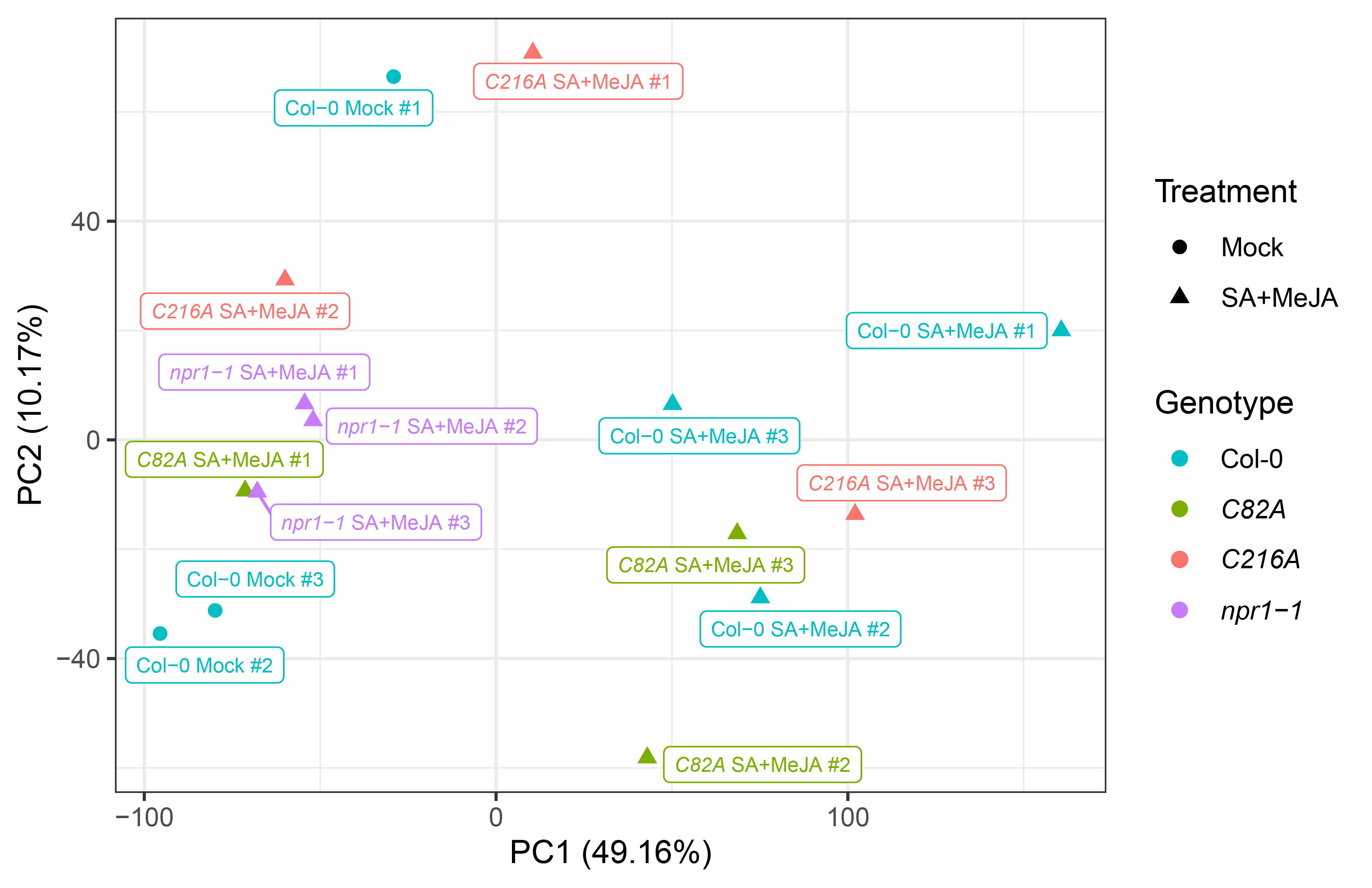

**Fig. S4** Principal component analysis of Mock-treated Col-0 and SA + MeJA-treated Col-0, *C82A*, *C216A*, and *npr1-1*

Principal component analysis was performed based on the log_2_ of the normalized read counts of all genes. Sample names indicate genotype, treatment and replicate number. Samples are colored according to their genotype. PC: principal component. # Indicates replicate number, corresponding to Fig. S3.

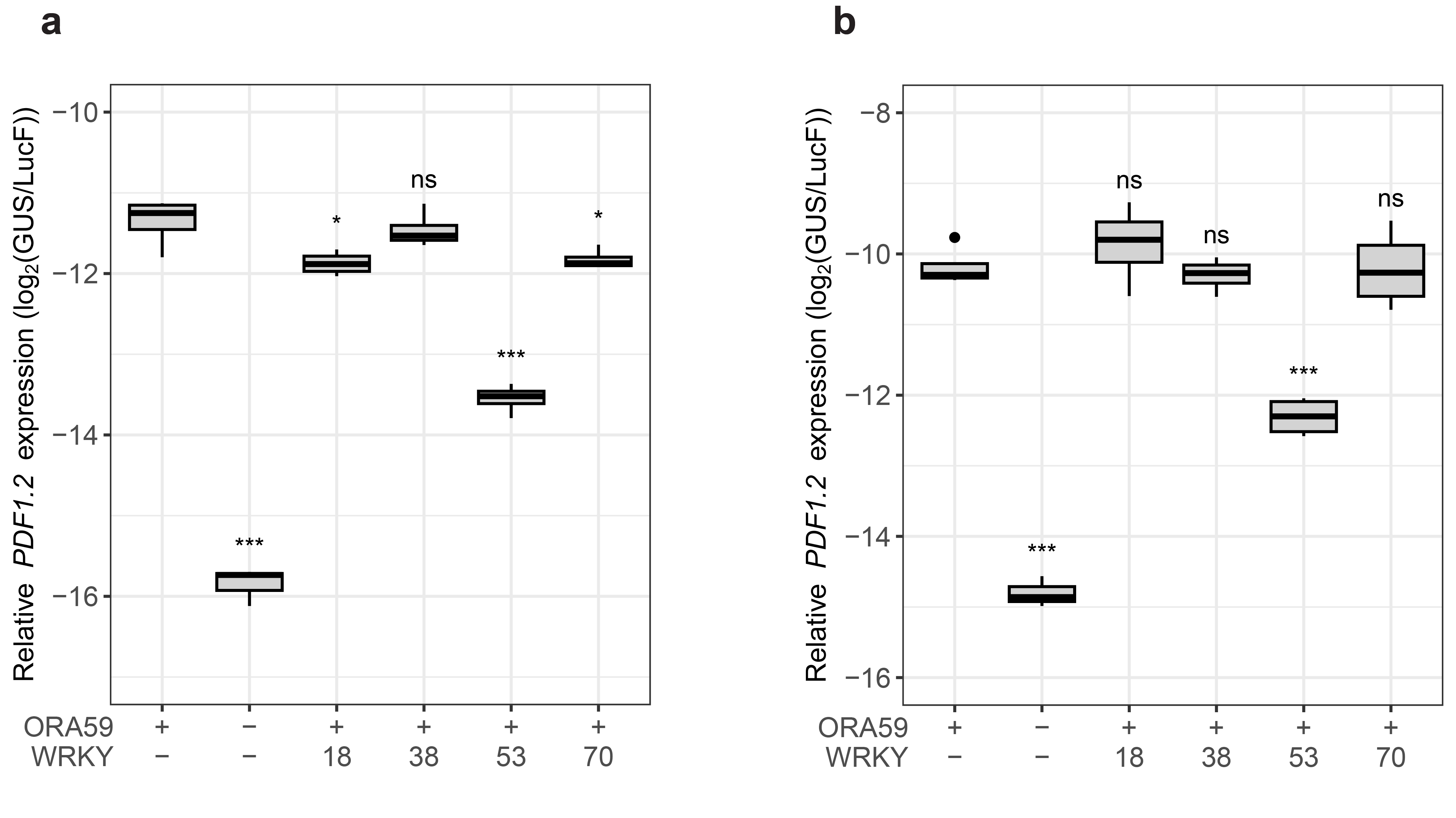

**Fig. S5** Suppression of ORA59-induced *PDF1.2* expression by four candidate WRKY TFs in Arabidopsis protoplasts

Protoplasts isolated from 3-week-old plants were transfected with a single plasmid described in (the legend of) Fig. 4a. Relative *PDF1.2* expression was determined as the log_2_ of relative GUS levels divided by relative LucF levels, as quantified using the substrates MUG and luciferin, respectively. Asterisks indicate significant differences from the reference (ORA59+ WRKY-) according to Dunnett's multiple-comparison test (*P < 0.05, **P < 0.01, ***P < 0.001, ns = not significant; *n*=4). Points that lie beyond 1.5 times the interquartile range are shown as dots in the plot, and were taken along with the other points in the statistical analysis. Panels **a** and **b** are two repeats of the same experiment. The third repeat is included as Fig. 4.

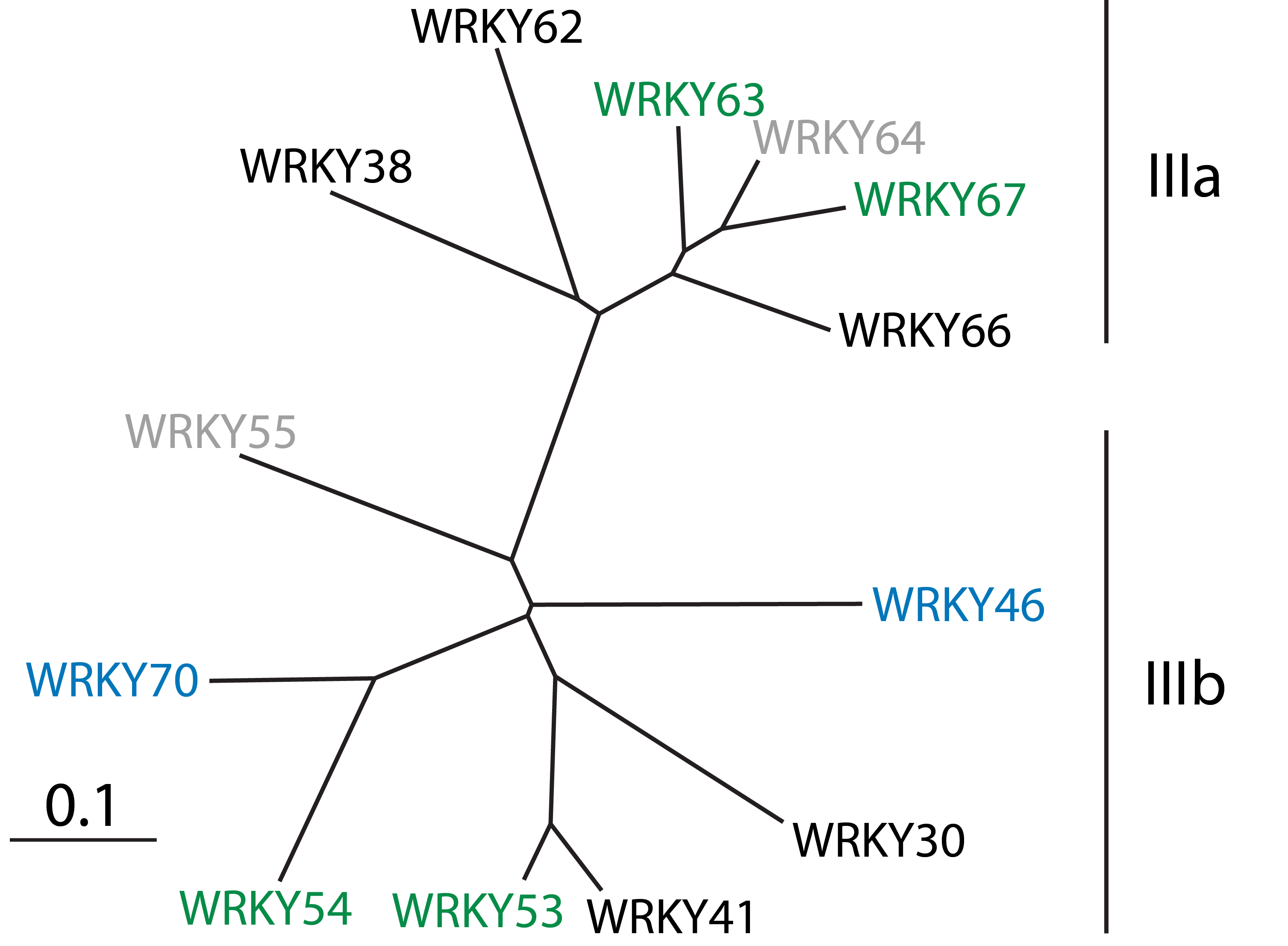

**Fig. S6** Phylogenetic analysis of the WKRY group III

Figure is adapted from Kalde et al. (2003). Phylogeny is based on the WRKY domain of each TF. The diagram shows an unrooted tree constructed by Kalde et al. (2003) using TREEVIEW (Page 1996). All WKRY TFs shown were tested for suppression of ORA59-induced *PDF1.2* expression in our protoplast expression assays, except WRKY55 and WRKY64, which we were unable to clone (indicated in gray). WRKY TFs that were consistently found to suppress ORA59-induced *PDF1.2* expression in our experiments are indicated in green. WRKY TFs that have been linked to SA/JA crosstalk in other papers, but were not (consistently) found to suppress ORA59-induced *PDF1.2* expression in our setup are indicated in blue. WRKY50, which we found suppressed ORA59-induced *PDF1.2* expression, is not included here because it does not belong to group III.

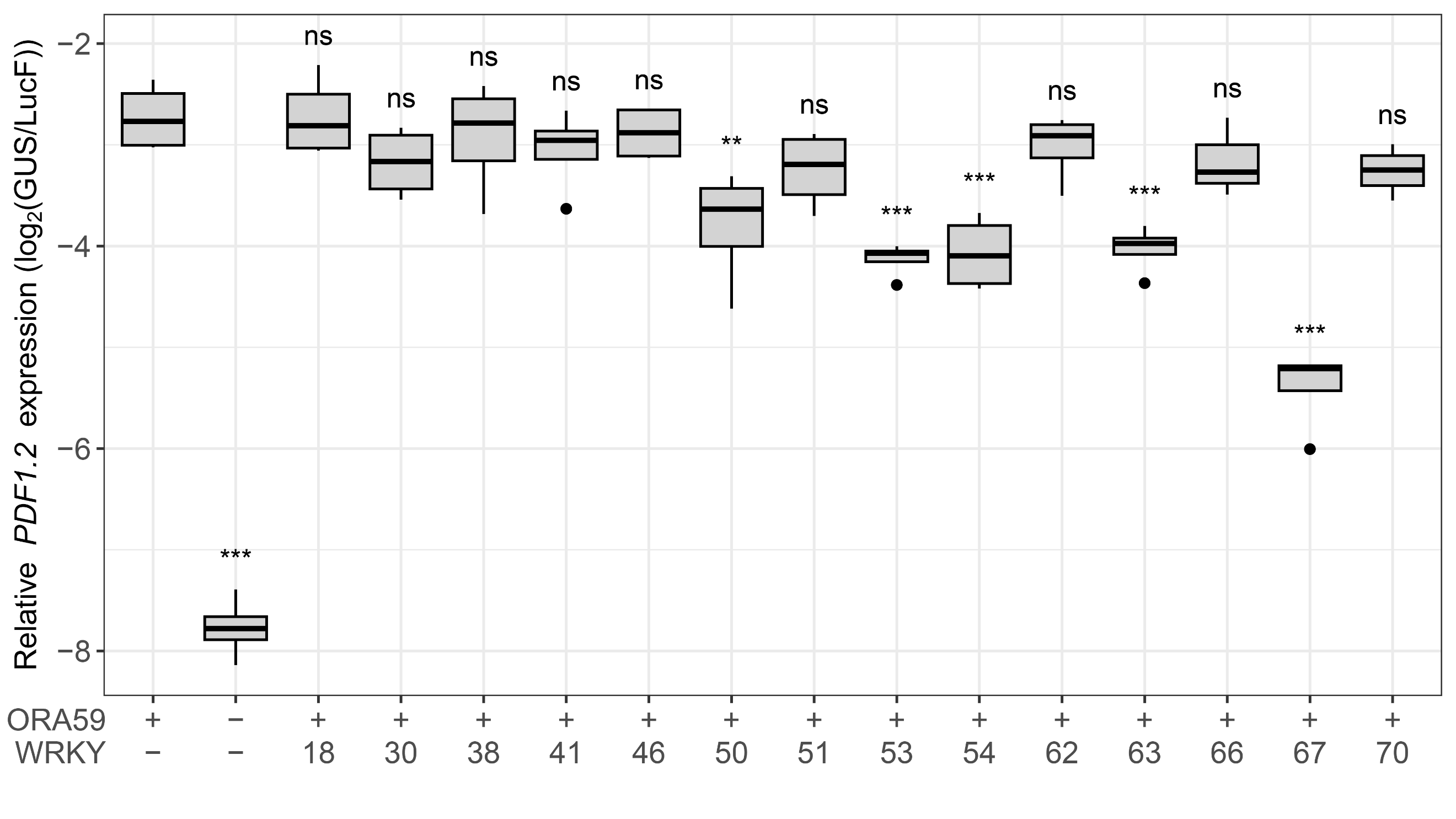

**Fig. S7** Suppression of ORA59-induced *PDF1.2* expression by selected WRKY TFs in Arabidopsis protoplasts

Protoplasts isolated from 3-week-old plants were transfected with a single plasmid, depicted and described in (the legend of) Fig. 4a. Relative *PDF1.2* expression was determined as the log_2_ of relative GUS levels divided by relative LucF levels, as quantified using the substrates MUG and luciferin, respectively. Asterisks indicate significant differences from the reference (ORA59+ WRKY-) according to Dunnett's multiple-comparison test (*P < 0.05, **P < 0.01, ***P < 0.001, ns = not significant; *n*=4). Points that lie beyond 1.5 times the interquartile range are shown as dots in the plot, and were taken along with the other points in the statistical analysis. A repeat of this experiment is depicted in Fig. 5.

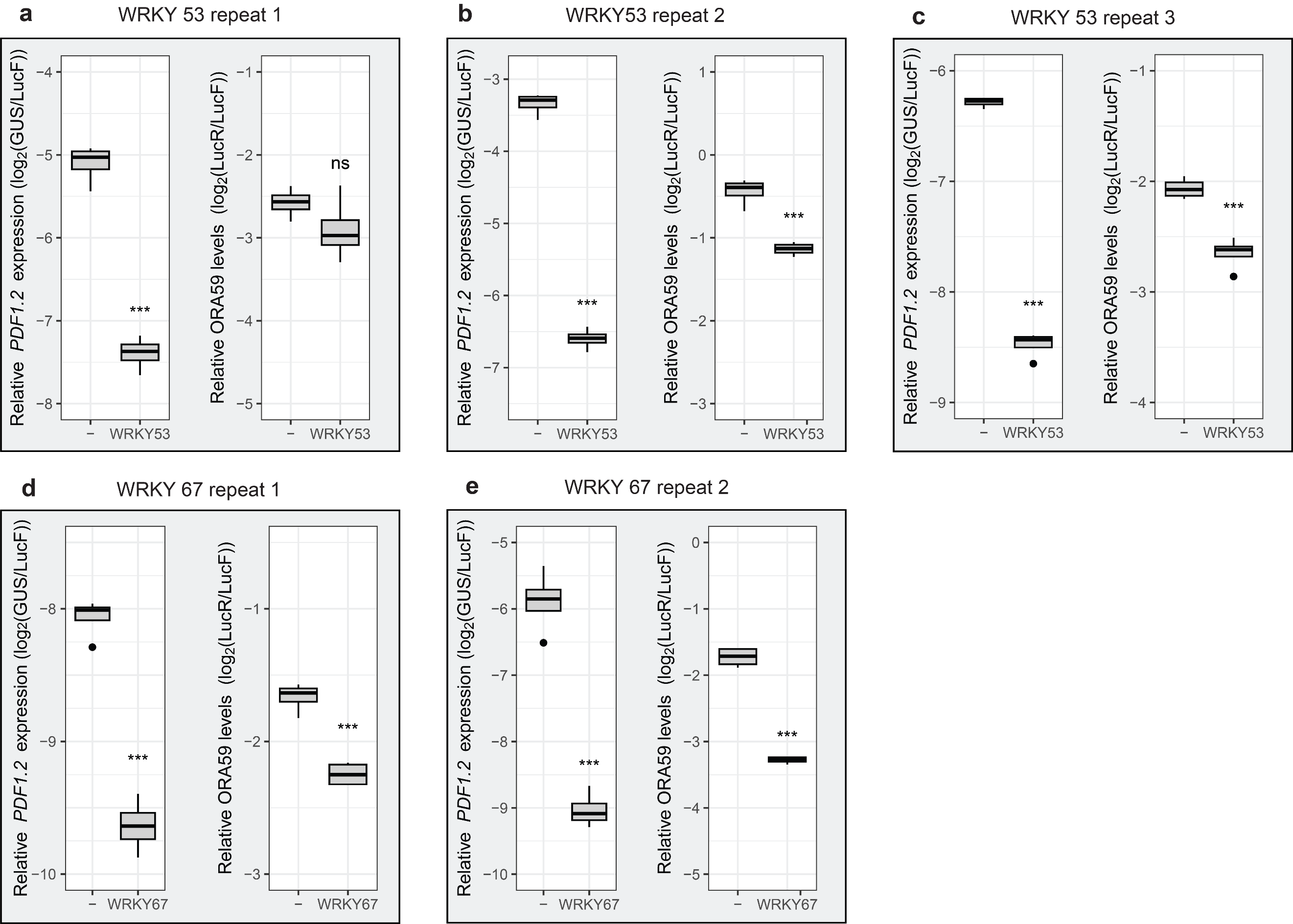

**Fig. S8** Suppression of *PDF1.2* expression and relative ORA59 protein levels by selected WRKY TFs

Protoplasts isolated from 3-week-old plants were transfected with two plasmids, described in (the legend of) Fig. 6a. In all panels, the left figure shows *PDF1.2* expression, which was determined as the log_2_ of relative GUS levels divided by relative LucF levels, as quantified using the substrates MUG and luciferin, respectively. The right figure shows relative ORA59 protein levels, which were determined as the log_2_ of relative LucR levels divided by relative LucF levels, as quantified using the substrates coelenterazine and luciferin, respectively. (**a-c**) three repeats of the same experiment with a plasmid containing either only the *35S_pro_:PDF1.2*, *35S_pro_:LucF* and *35S_pro_:ORA59-LucR* modules (“-“) or the same plasmid co-transfected with a plasmid containing *35S_pro_:WRKY53* (“WRKY53”). (**d, e)** two repeats of the same experiment with a plasmid containing either only the *35S_pro_:PDF1.2*, *35S_pro_:LucF* and *35S_pro_:ORA59-LucR* modules (“-“) or the same plasmid co-transfected with a plasmid containing *35S_pro_:WRKY67* (“WRKY67”). Asterisks significant differences from the reference (-) according to Dunnett's multiple-comparison test (*P < 0.05, **P < 0.01, ***P < 0.001, ns = not significant; *n*=4). Points that lie beyond 1.5 times the interquartile range are shown as dots in the plot, and were taken along with the other points in the statistical analysis. Fig. 6 shows a similar experiment where WRKY50, WRKY53, WRKY54, WRKY63 and WRKY67 were tested.

**SUPPLEMENTARY TABLES**

**Table S1** List of primers used for qRT-PCR in this study

Primer name and sequence (5’ to 3’) are denoted.

| **Name** | **Sequence (5ˈ-> 3ˈ)** |
| --- | --- |
| AT1G13320_fw | TAACGTGGCCAAAATGATGC |
| AT1G13320_rv | GTTCTCCACAACCGCTTGGT |
| PR1_fw | CTCGGAGCTACGCAGAACAACT |
| PR1_rv | TTCTCGCTAACCCACATGTTCA |
| PDF1.2_fw | TTTGCTGCTTTCGACGCAC |
| PDF1.2_rv | CGCAAACCCCTGACCATG |
| VSP2_fw | ACGGAACAGAGAAGACCGAC |
| VSP2_rv | TCTTCCACAACTTCCAACGG |

**Table S2** Primers used for amplification of level 0 plasmids and vector name after level 0 reaction

The part of the primer designed to overlap with the DNA template is denoted in uppercase and any added nucleotides are denoted in lowercase (except in the EM primer, which does not match any piece of existing DNA). The four base pairs that form the overhang after the BbsI restriction reaction are denoted in blue. In cases where a nucleotide was altered in the overhang to take out an internal restriction site, this nucleotide is denoted in red. The restriction recognition site is denoted in green. The fragments were cloned into the plasmid denoted under ‘New plasmid name’ before the underscore using a Golden Gate reaction with BbsI. The ‘New plasmid name’ is the name used for the plasmid after cloning.

| **Name** | **Sequence** | **Fragment size (bp)** | **New plasmid name** |
| --- | --- | --- | --- |
| PDF1.2_BbsI_fw | gatcgaagacatggagGCATGCATCGCCGCATCGATATCC | 1230 | pICH41295_PDF1.2 |
| PDF1.2_BbsI_rv | gactgaagacatcattGATGATTATTACTATTTTGTTTTCAATGTATAGATGTATG |  |  |
| ORA59_ BbsI_1_fw | gactgaagacataATGGAATATCAAACTAACTTCTTAAGTGGAGAGTTTTC | 632 | pAGM1287_ORA59 (no stop codon version)  pICH41308_ORA59  (stop codon version) |
| ORA59_ BbsI_1_rv | gactgaagacagAgGACGAAGAAGATGAATAGGAGGAGGAG |  |  |
| ORA59_ BbsI_2_fw | gactgaagaccgTCcTCTTCTTCTTTGTCGTCAAGAAGTAGAAAACAGAG | 159 |  |
| ORA59_ns_ BbsI_2_rv^a^ | gactgaagacatcgaaccAGAACATGATCTCATAAGCTCTTCTAAGTATTC |  |  |
| ORA59_stop_ BbsI_2_rv^b^ | gactgaagacataagcTCAAGAACATGATCTCATAAGCTCTTCTAAGTATTC | 160 |  |
| WRKY18_BbsI_1_fw | gactgaagacataATGGACGGTTCTTCGTTTCTCGACATC | 108 | pICH41308_WRKY18 |
| WRKY18_BbsI_1_rv | gactgaagactaGAaACCTCCTTCTTCGGAAGTTTTGCGG |  |  |
| WRKY18_BbsI_2_fw | gactgaagacatTtTCAGTTTTGGCTTCTACTCACTTAAAGAGG | 882 |  |
| WRKY18_BbsI_2_rv | gactgaagacataagcTCATGTTCTAGATTGCTCCATTAACCTCCC |  |  |
| WRKY30_BbsI_fw | gactgaagacataATGGAGAAGAACCATAGTAGTGGA | 941 | pICH41308_WRKY30 |
| WRKY30_BbsI_rv | gactgaagacataagcCTAAGAATAGAACCCACCAAATCC |  |  |
| WRKY38_BbsI_fw | gactgaagactaaATGGAAATGAACTCCCCACACGAAAAGG | 870 | pICH41308_WRKY38 |
| WRKY38_BbsI_rv | gactgaagacataagcTCAAAAGTAAAACTGATCATAACGATCCCACG |  |  |
| WRKY41_BbsI_1_fw | gactgaagactaaATGAATGGAAATGATGAATTGGGAGC | 113 | pICH41308_WRKY41 |
| WRKY41_BbsI_1_rv | gactgaagactaAtGACGAACCCTGAAGCTGC |  |  |
| WRKY41_BbsI_2_fw | gactgaagactaTCaTCTCCATCGTTGTCAGCTTC | 887 |  |
| WRKY41_BbsI_2_rv | gactgaagactaaagcTTAAATCGAATTGTGGAAAAAAGTGG |  |  |
| WRKY46_BbsI_1_fw | gactgaagacctaATGATGATGGAGGAGAAACTTGTGAATCAACGAA | 254 | pICH41308_WRKY46 |
| WRKY46_BbsI_1_rv | gactgaagacctAaACCTTCTTAGAATCCTTTCCATCGATCTCAAGGCT |  |  |
| WRKY46_BbsI_2_fw | gactgaagacaaGTtTTCAAAAAGAGGAAAGTATCGGA | 310 |  |
| WRKY46_BbsI_2_rv | gactgaagacaaTtTTCGGGGATGTGATGTTGTT |  |  |
| WRKY46_BbsI_3_fw | gactgaagacaaAAaACGACGACGAACTTCTCTGTT | 110 |  |
| WRKY46_BbsI_3_rv | gactgaagacaaCcTCCGATTGCTCTGTAACATG |  |  |
| WRKY46_BbsI_4_fw | gactgaagacaaGAgGACATGAAACCGACCAAGTC | 328 |  |
| WRKY46_BbsI_4_rv | gactgaagacaaaagcCTACGACCACAACCAATCCTG |  |  |
| WRKY50_BbsI_fw | gactgaagacataATGAATGATGCAGACACAAACTT | 551 | pICH41308_WRKY50 |
| WRKY50_BbsI_rv | gactgaagactaaagcCTAGTTCATGCTAGAGTGATTGTGGGAACCCTC |  |  |
| WRKY51_BbsI_fw | gactgaagacataATGAATATCTCTCAAAACCCTAGCC | 514 | pICH41308_WRKY51 |
| WRKY51_BbsI_rv | GactgaagactaaagcTTAAGATCGAAGAAGAGAGTGTTGG |  |  |
| WRKY53_BbsI_fw | gactgaagacaaaATGGAAGGAAGAGATATGTTAAGTTGG | 1004 | pICH41308_WRKY53 |
| WRKY53_BbsI_rv | gactgaagacaaaagcTTAATAATAAATCGACTCGTGTAAAAAC |  |  |
| WRKY54_BbsI_1_fw | gactgaagacataATGGATTCGAATAGTAACAACACG | 161 | pICH41308_WRKY54 |
| WRKY54_BbsI_1_rv | gactgaagacatTaTCATCGATGTGGTACTGGTTAGA |  |  |
| WRKY54_BbsI_2_fw | gactgaagacatGAtACCCGTCTTGTTTCCGG | 352 |  |
| WRKY54_BbsI_2_rv | gactgaagacatCcTCAGAACTTTTAGCTTCCACGA |  |  |
| WRKY54_BbsI_3_fw | gactgaagacatGAgGACAGATATGCTTGGAGGAAAT | 235 |  |
| WRKY54_BbsI_3_rv | gactgaagacatTtTTCGCGTGCGTTTGGT |  |  |
| WRKY54_BbsI_4_fw | gactgaagacatAAaACCGAGCCTTTTGATCAAGA | 412 |  |
| WRKY54_BbsI_4_rv | gactgaagacataagcTCACATAGCACTTGTTCTTTCATAA |  |  |
| WRKY55_BbsI_fw | gactgaagacataATGTACTCGTACAAAAAAATAAGTTACCA | 911 | Unable to obtain a correct plasmid |
| WRKY55_BbsI_rv | gactgaagacataagcCTAAGATGGGTCGTTCTTTGGAAATATGAAATCCATGTTGGTTC |  |  |
| WRKY62_BbsI_1_fw | gactgaagacataATGAACTCTTGCCAACAAAAGG | 720 | pICH41308_WRKY62 |
| WRKY62_BbsI_1_rv | gactgaagacatGtGACCAATCTTCTGCTATCACCG |  |  |
| WRKY62_BbsI_2_fw | gactgaagacatTCaCCATCCCAGTGGATGTCG | 120 |  |
| WRKY62_BbsI_2_rv | gactgaagacataagcTCATGATGATAAGTCGTGAGATG |  |  |
| WRKY63_BbsI_1_fw | gactgaagacataATGTTTTCAAACATCGATCACAAGG | 225 | pICH41308_WRKY63 |
| WRKY63_BbsI_1_rv | gactgaagacatAtGACTCATGGTGTGGTTGACG |  |  |
| WRKY63_BbsI_2_fw | gactgaagacatTCaTCTTCTCGAGACATGGCAGG | 553 |  |
| WRKY63_BbsI_2_rv | gactgaagacatCGATGAAAATAGAGGAAATTCATTCCC |  |  |
| WRKY63_ann_3_fw^c^ | ATCG GAgGACCTGATGTTGTTTTGA | 29 |  |
| WRKY63_ann_3_rv^c^ | aagc TCAAAACAACATCAGGTCcTC |  |  |
| WRKY64_BbsI_1_fw | gactgaagacataATGTTCTCCAACATCGATCAAA | 200 | Unable to obtain a correct plasmid |
| WRKY64_BbsI_1_rv | gactgaagacatTAGGCGGGTCCACAAAAG |  |  |
| WRKY64_ann_2_fw^c^ | CCTAGcCTTCTACCACACAATGAGTCaTC | 33 |  |
| WRKY64_ann_2_rv^c^ | AAGAGAtGACTCATTGTGTGGTAGAAGgC |  |  |
| WRKY64_BbsI_3_fw | gactgaagacatTCTTCAAAACATGACAAGTCATGTTC | 536 |  |
| WRKY64_BbsI_3_rv | gactgaagacatCcTCCGATGAAAATGGAGGAAATTG |  |  |
| WRKY64_ann_4_fw^c^ | GAgGACATTATGTTTTTTGACAATATTGCTAATCTTGATTAG | 46 |  |
| WRKY64_ann_4_rv^c^ | aagcCTAATCAAGATTAGCAATATTGTCAAAAAACATAATGT |  |  |
| WRKY66_BbsI_1_fw | gactgaagacataATGTCTCTTGAGATTGATGCGAAG | 195 | pICH41308_WRKY66 |
| WRKY66_BbsI_1_rv | gactgaagacatAtGACAGAGCAAACGAGAAAGAATC |  |  |
| WRKY66_BbsI_2_fw | gactgaagacatTCaTCTCAGAACATACCACGTCATG | 540 |  |
| WRKY66_BbsI_2_rv | gactgaagacataagcTTAAGATTTATTAATGTTCAATCCTGGAAAG |  |  |
| WRKY67_BbsI_fw | gactgaagacataATGGTTTCCAACATTGATCACAAG | 794 | pICH41308_WRKY67 |
| WRKY67_BbsI_rv | gactgaagacataagcCTAATCAAAAGCAGAAATGTTTTCCAAG |  |  |
| WRKY70_BbsI_1_fw | gactgaagacataATGGATACTAATAAAGCAAAAAAGCTTAAAGTTATGAACC | 268 | pICH41308_WRKY70 |
| WRKY70_BbsI_1_rv | gactgaagactaCcTCTTCCTTCATTGAGGTAGATAAACTAGGAC |  |  |
| WRKY70_BbsI_2_fw | gactgaagacgaGAgGACAATCCTCATCGTCATCATGGTTCGTC | 732 |  |
| WRKY70_BbsI_2_rv | gactgaagacataagcTCAAGATAGATTCGAACATGAACTGAAGATAGAG |  |  |
| LucR_BbsI_1_fw | gatcgaagacatttcgATGGCTTCGAAAGTTTATGATCC | 879 | pAGM1301_LucR |
| LucR_BbsI_1_rv | gatcgaagacattAGACCTTTTACTTTGACAAATTCAGTATTAGG |  |  |
| LucR_BbsI_2_fw | gatcgaagacatTCTaCATTTTTCGCAAGAAGATGC | 125 |  |
| LucR_BbsI_2_rv | gatcgaagacataagcCTAGATAGATCTTTGTTCATTTTTGAGAA |  |  |
| EM_ann_fw^c^ | AATGTGATCGAGAATTCGAGCTCGAGCTCGGTAC | 34 | pICH41308_EM |
| EM_ann_rv^c^ | AAGCGTACCGAGCTCGAGCTCGAATTCTCGATCA |  |  |

^a^This primer does not include the stop codon.

^b^This primer includes the stop codon.

^c^This primer was used for an annealing-based protocol rather than PCR.

**Table S3** DNA components for assembly of level 1 plasmids for transient protoplast assays

Components were assembled using Golden Gate cloning with BsaI. Plasmids were derived from level 0 reactions (Table S2) or the MoClo plant parts toolkit (Weber et al. 2011; Werner et al. 2012). Name codes represent level, intended position in the level 2 vector and a unique number, all separated by dots. NB: the unique number does not start with 1 because the plasmids were part of a series of experiments with different plasmids.

| **Description** | **Backbone vector** | **Promoter + 5ˈ UTR** | **CDS** | **3ˈ tag** | **3ˈ UTR + terminator** | **New vector name** |
| --- | --- | --- | --- | --- | --- | --- |
| *PDF1.2_pro_:GUS* | pICH47732 | pICH41295_PDF1.2 | pICH75111 | - | pICH41414 | 1.1.2 |
| *35S_pro_:LucF* | pICH47742 | pICH51277 | pICSL80001 | - | pICH41414 | 1.2.2 |
| *35S_pro_:ORA59* | pICH47751 | pICH51277 | pICH41308_ORA59 | - | pICH41414 | 1.3.1 |
| *Empty module position 3* | pICH47751 | pICH51277 | pICH41308_EM | - | pICH41414 | 1.3.2 |
| *35S_pro_:ORA59-LucR* | pICH47751 | pICH51277 | pAGM1287_ORA59 | pAGM1301_LucR | pICH41414 | 1.3.3 |
| *35S_pro_:WRKYxx^a^* | pICH47751 | pICH51277 | pICH41308_WRKYxx^a^ | - | pICH41414 | 1.4.xb |
| *Empty module position 4* | pICH47751 | pICH51277 | pICH41308_EM | - | pICH41414 | 1.4.5 |
| *35S_pro_:EYFP* | pICH47772 | pICH51277 | pICSL80014 | - | pICH41414 | 1.5.1 |

^a^xx refers to a unique WRKY TF number.

bx refers to a unique number for each WRKY TF level 1 construct (except 5 which refers to the construct with an empty module instead of a WRKY TF).

**Table S4** DNA components for assembly of level 2 plasmids for a transient protoplast assay

Plasmids were derived from level 1 reactions (Table S3) or the MoClo plant parts toolkit (Weber et al. 2011; Werner et al. 2012). All constructs had pAGM4673 as acceptor vector and used end-linker 5 to link the fragment of position 5 to the backbone vector.

| **Construct description** | **Position 1** | **Position 2** | **Position 3** | **Position 4** | **Position 5** |
| --- | --- | --- | --- | --- | --- |
| Control plasmid for *PDF1.2* induction | 1.1.2 | 1.2.2 | 1.3.2 | 1.4.5 | 1.5.1 |
| Control plasmid for WRKY repression | 1.1.2 | 1.2.2 | 1.3.1 | 1.4.5 | 1.5.1 |
| Plasmid including WRKY TF | 1.1.2 | 1.2.2 | 1.3.1 | 1.4.x^a^ | 1.5.1 |
| Readout plasmid for ORA59 degradation (transfected together with level 1 WRKY-containing plasmid) | 1.1.2 | 1.2.2 | 1.3.3 | pICH54044 | 1.5.1 |
| Plasmid for ORA59 degradation including WRKY TF | 1.1.2 | 1.2.2 | 1.3.3 | 1.4.x^1^ | 1.5.1 |

^a^x refers to any number of a construct that contains a WRKY TF.
